# Inherited CDA-I disease: anemia-associated mutations disrupt CDIN1-Codanin1 complex

**DOI:** 10.1101/2023.05.25.542057

**Authors:** Martin Stojaspal, Tomáš Brom, Ivona Nečasová, Tomáš Janovič, Pavel Veverka, Naina Verma, Lukáš Uhrík, Lenka Hernychova, Ctirad Hofr

## Abstract

Congenital dyserythropoietic anemia type I (CDA-I) is a rare hereditary disease marked by ineffective erythropoiesis, a characteristic spongy heterochromatin structure in erythroblasts, and mutations in the genes *CDAN1* and *CDIN1,* which encode the proteins Codanin1 and CDIN1. Codanin1 regulates histone shuttling via the chaperone ASF1, yet the role of CDIN1 in CDA-I pathology remains unclear. Notably, CDIN1 is known to interact directly with the C-terminus of Codanin1. Although mutations in both genes are critical to the disease phenotype, their molecular-level effects have not been fully elucidated. Here, we present a comprehensive structural and functional analysis of the CDIN1-Codanin1 C-terminus complex. Using complementary biophysical techniques, we show that CDIN1 and Codanin1 C-terminus form a high-affinity heterodimeric complex with equimolar stoichiometry. We further delineate the essential interacting regions of CDIN1 and Codanin1. We demonstrate that CDA-I-associated mutations in either protein disrupt CDIN1-Codanin1 interaction, suggesting a potential molecular mechanism underlying the disease.

**Graphical Abstract:** 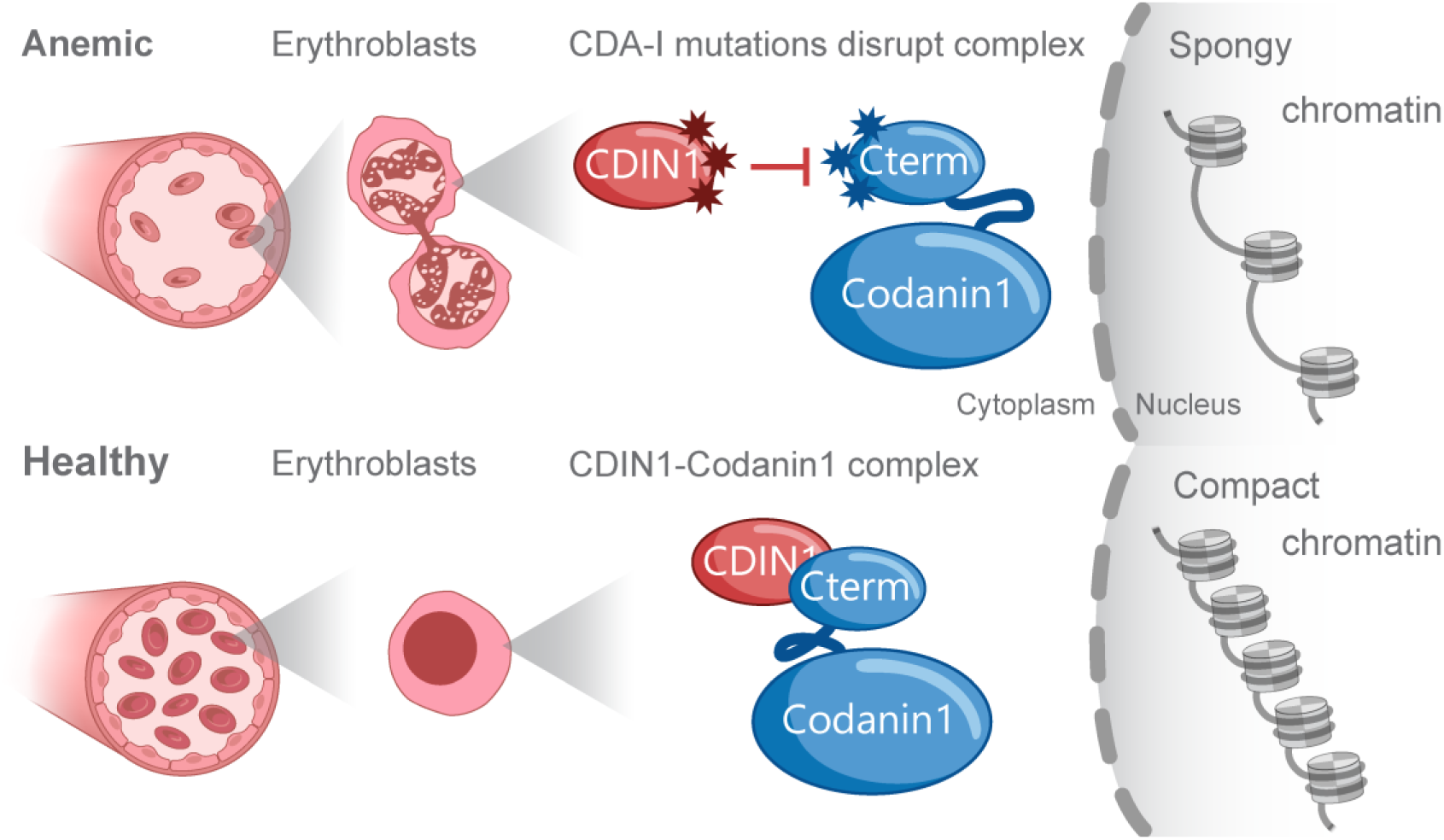

## Introduction

Congenital dyserythropoietic anemia (CDA) is a rare hereditary disease manifesting abnormal erythroblast morphology and ineffective erythropoiesis in the bone marrow^1^. CDA type I (CDA-I) incidence is 1 per ∼ 207,000 live births, according to a recent allele frequency analysis^2^. CDA- I causes spleen enlargement and iron overload. At the cytological level, CDA-I is associated with increased mean red blood cell volume in 75% of all erythroblasts. Additionally, CDA-I induces binucleate erythroblasts (up to 7%) and internuclear chromatin bridges (up to 3%). Most strikingly, electron microscopy revealed spongy heterochromatin with a “Swiss-cheese” like appearance in up to 60% of cells^3^.

From a genetic point of view, CDA-I is an autosomal recessive disease caused by biallelic mutations in genes *CDAN1* and *CDIN1* (originally called C15orf41), which account for approximately 90% of all CDA-I cases^4^. *CDAN1* was first identified as the causative gene in a cohort of Israeli Bedouins^5^. *CDAN1* is essential for cell survival as mice embryos with an artificially disrupted gene perished in the early stages of development^6^. The highly conserved *CDAN1* gene^5^ encodes Codanin1 (Fig. 1A), a 134 kDa protein that negatively regulates the shuttling of newly synthesized histones from the cytoplasm to the nucleus by histone chaperone ASF1 (anti-silencing function protein 1)^7^. The second causative gene – *CDIN1*, was identified by whole-genome sequencing of individuals from CDA-I disease-affected pedigrees^8^. *CDIN1* encodes the 32 kDa CDAN1-interacting nuclease 1 (CDIN1) of unknown function (Fig. 1B). Although CDIN1 was initially predicted as a restriction nuclease^8^, no evidence of nuclease activity has been published yet. Gene expression arrays revealed that *CDIN1* is most extensively transcribed in hematopoietic stem cells, B lymphoblasts, cardiomyocytes, and fetal liver, suggesting the essentiality of CDIN1 in hematopoiesis^9^.

**Fig. 1:**
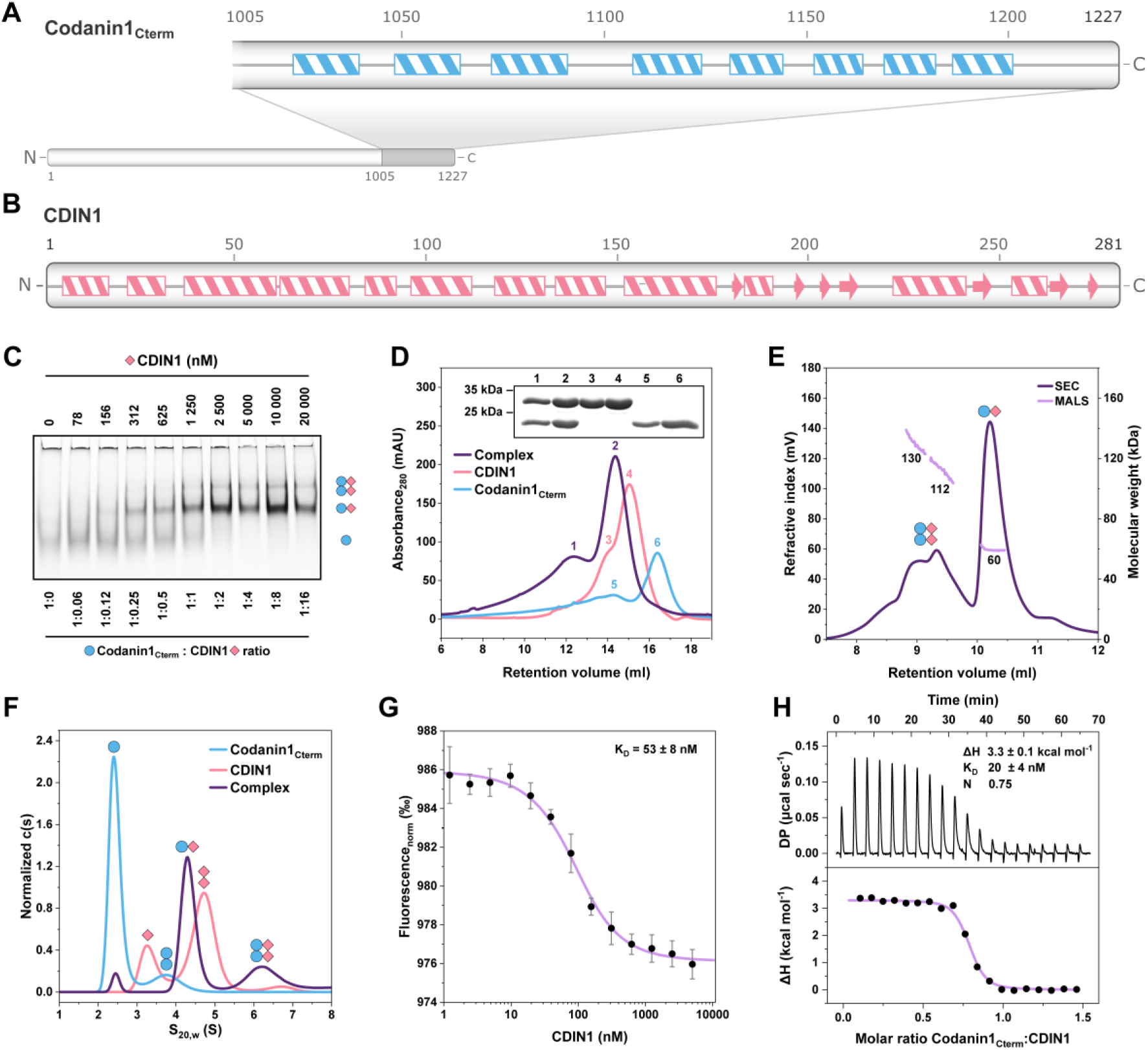
CDIN1 and Codanin1_Cterm_ – structure, binding stoichiometry and affinity. (A) Codanin1_Cterm_ and (B) CDIN1 size and secondary structure according to AlphaFold prediction; boxes denominate alpha helices, and arrows represent beta sheets. (C) EMSA shows the binding of CDIN1 to fluorescently labeled Codanin1_Cterm_ with highlighted molar ratios. Suggested protein arrangements are depicted as pictograms at band levels. (D) Size-exclusion chromatograms of individual and equimolarly incubated proteins demonstrate that CDIN1 and Codanin1_Cterm_ form stable complexes. SDS-PAGE analysis of protein content in corresponding SEC fractions (inset). (E) SEC-MALS of CDIN1-Codanin1_Cterm_ complex cross-validates heterodimer formation with corresponding molecular weight. (F) Non-overlapping AUC profiles for individual proteins and complexes show heterodimer formation. (G) MST reveals the nanomolar affinity of CDIN1 binding to Codanin1_Cterm_. Data points are associated with error bars representing the standard deviation calculated from five independent measurements. (H) ITC titration of Codanin1_Cterm_ (110 μM) to CDIN1 (11 μM) also supports high mutual binding affinity at micromolar protein concentration.

Previous co-immunoprecipitation and immunofluorescence studies performed on transfected mammalian cell lines HEK293T, HeLa, and U2OS have shown that CDIN1 interacts directly with the C-terminal region of Codanin1^2,4,10^. Swickley *et al.* and Shroff *et al*. independently found that CDIN1 binds the C-terminal part of Codanin1 (Codanin1_Cterm_) comprising amino acids 1005-1227^4,10^. However, interacting regions of CDIN1 and Codanin1 have not been mapped and precisely defined.

The most recent genetic testing and molecular diagnoses revealed eight missense mutations in Codanin1_Cterm_ and six mutated amino acids in CDIN1 related to CDA-I disease progression^2,11^.

To date, pathological CDA-I mutations have not been linked to CDIN1-Codanin1 interaction. As effects of individual mutations remain unclear, direct causative links connecting specific mutations and chromatin changes are still missing. Additionally, while the significance of CDIN1 and Codanin1 in CDA-I pathology is recognized, it is still unclear if the newly identified protein CDIN1 plays a part in histone trafficking, or how exactly CDIN1 influences Codanin1’s other functions and the pathology of CDA-I.

In this study, we demonstrate that CDIN1 and Codanin1_Cterm_ bind with a high affinity. We present small-angle X-ray scattering data of CDIN1 and Codanin1_Cterm_, along with their stable complex and describe binding stoichiometry. We further employed hydrogen-deuterium exchange mass spectrometry (HDX-MS) to identify the interacting regions that are critical for the binding of CDIN1 and Codanin1. Finally, we show that the CDA-I-associated mutations in interacting regions disturb CDIN1-Codanin1 complex. The revealed direct impact of disease-related mutations in CDIN1 and Codanin1 is essential for determining the molecular mechanism of their mutual interaction and identifying their roles in regulating histone reshuffling and abnormal heterochromatin arrangements. The new findings provide foundations for developing targeted biological therapies of CDA-I.

## Methods

### Origin of CDIN1 and Codanin1 coding sequences

The total RNA was extracted from 3 x 10^6^ HEK293T cells using the RNeasy Mini Kit (QIAGEN). RNA served as a template for subsequent reverse transcription into cDNA by iScript Reverse Transcription Supermix for RT qPCR (Bio-Rad). Specific primers (cDNA_CDIN1_Fw ATGATACTGACCAAAGCTCAGTAC and cDNA_CDIN1_Rv TCAAGCTATGCTGTGGCATAA) were designed to amplify CDIN1 coding sequence using proofreading Q5 High Fidelity DNA Polymerase (New England Biolabs). The PCR product was purified by GenElute PCR Clean Up Kit (Sigma Aldrich) and cloned into pCR Blunt II TOPO vector by Zero Blunt TOPO PCR Cloning Kit (Invitrogen). All procedures were done according to the manufacturer’s protocols. Plasmids containing the whole human Codanin1 sequence Topo_pCR2_CDAN1 and Codanin1 sequence optimized for bacterial expression pHAT4_CDAN1 were kindly provided by Prof. Anja Groth (University of Copenhagen).

### Cloning, expression, and purification of CDIN1 and Codanin1 variants

NEBuilder® HiFi DNA Assembly kit (New England Biolabs) was used to clone CDIN1 and codon-optimized Codanin1_Cterm_ coding sequences into HindIII, XhoI restriction site of plasmid pl21 (TriAltus) that is suitable for Im7/CL7 ultra-high affinity chromatography purification from bacteria. Similarly, Flag- and Myc-tagged variants of CDIN1, Codanin1, and Codanin1_Cterm_ were cloned into NotI, SalI restriction site of plasmid pHAGE_puro (Addgene #118692) that allows expression in mammalian cells. Cloning was performed according to the manufacturer’s recommendation. All constructs were transformed into *E. coli* strain One Shot™ TOP10 (Invitrogen) for plasmid amplification or BL21-CodonPlus (DE3)-RIPL Competent Cells (Agilent) for protein expression in bacteria. Successful cloning was confirmed by Sanger sequencing.

To express protein genes, BL21(DE3) RIPL cell cultures harboring pl21_CDIN1 and pl21_Codanin1_Cterm_ were grown in Terrific Broth (TB) medium containing 50 μg ml^-1^ kanamycin and 34 μg ml^-1^ chloramphenicol in orbital shaker New Brunswick™ Innova® 43R (New Brunswick) at 160 rpm and 37 °C. When OD_600_ reached a value between 0.6-0.8, the cell cultures were cooled down to 20 °C, and protein expression was induced by IPTG addition to the final concentration of 0.1 mM. After induction, the cell cultures were cultured O/N and collected the next day by centrifugation (8 000 g, 8 min, 4 °C) and stored at -80 °C.

The pellet was resuspended in lysis buffer (20 mM Tris-HCl, 1 000 mM NaCl, 5% glycerol, 0.05% Triton X-100, pH 8.0 with the addition of 1 mM DTT, protease inhibitor cocktail cOmplete tablets EDTAfree (Roche), and 0.4 mg ml^-1^ of lysozyme before use), using 5 ml of lysis buffer per gram of wet pellet. The cell suspension was sonicated on ice with amplitude settings 30 for 4 min of process time, where the pulseon and pulseoff intervals were 1 and 3 seconds, respectively (Misonix S-4000). The cell lysate was centrifugated at 60 000 g, 4 °C for 30 minutes and filtered through 0.45 μm Sterivex™ (Merck Millipore). Supernatants of CDIN1 and Codanin1_Cterm_ were purified by a three-step protocol utilizing CLiM™ CL7/Im7 Affinity Tag System (TriAltus), HisTrap-HP (Cytiva), and size-exclusion chromatography (Cytiva) using flow rate 1 ml min^-1^ unless stated otherwise. Firstly, filtered supernatant was loaded onto the Im7 FPLC column (5 ml), and then the column was washed by five column volumes of Im7 buffer (20 mM Tris-HCl, 500 mM NaCl, 5% glycerol, pH 8.0). To elute protein, the proteolytic cleavage was done by homemade His-tagged Ulp1 protease diluted in Im7 buffer (final concentration 0.1 mg ml^-1^) with the addition of 1 mM DTT. To improve yield, the flow was paused, and the column was incubated three times for 15 minutes. An elution sample from Im7 FPLC column was loaded onto the HisTrap-HP column to clear the sample from residual His-tagged Ulp1. In the case of CDIN1 and Codanin1_Cterm_, non-specific binding to HisTrap-HP column was observed. To reduce non-specific binding and elute proteins, a 5-10% gradient of His-Trap buffer (20 mM Tris-HCl, 500 mM NaCl, 500 mM imidazole, 5% glycerol, pH 8.0) in Im7 buffer was used. For the buffer exchange and final exclusion of possible impurities, CDIN1 and Codanin1_Cterm_ were loaded onto the HiLoad 16/600 column containing Superdex 200 pg (Cytiva) and resolved using final buffer (20 mM Tris-HCl, 150 mM NaCl, 5% glycerol, pH 8.0) that was used in most experiments afterward. In the last step, proteins were concentrated by ultrafiltration on 10K Amicon® Ultra Centrifugal Filters (Merck Millipore) and either fluorescently labelled or immediately aliquoted, snap-freezed in liquid nitrogen, and stored at -80 °C. Protein identity was analyzed by MALDI-TOF mass spectrometry (Proteomics Core Facility, CEITEC MU, Brno). Protein purity was assessed by densitometry analysis of protein samples resolved on 12% SDS-PAGE gels stained with Coomassie Brilliant Blue G250^12^. Protein samples were thawed on ice prior to each experiment to maintain a uniform method across all the studies shown here.

### Protein labeling by fluorescent dyes

Fluorophores were covalently attached to proteins, allowing their visualization during interaction studies using fluorescence-based methods such as EMSA and MST, as described below. Proteins were labeled using Alexa Fluor 488 coupled with carboxylic acid, 2,3,5,6-tetrafluorophenyl ester (Invitrogen) or Alexa Fluor 594 coupled with carboxylic acid, succinimidyl ester (Invitrogen) in 4-fold molar excess over protein. First, fluorophores were diluted in 1 M sodium bicarbonate in one-tenth of the protein final volume, mixed with protein (0.5 mg), and incubated for 1 h at 4 °C while stirring. The mixture was loaded on gravity flow PD-10 columns (GE Healthcare) and isocratically eluted with 20 mM Tris-HCl, 150 mM NaCl, 5% glycerol, pH 8.0. The degrees of labeling (dye : protein ratio) over 90% were confirmed by UV/Vis spectroscopy.

### Electrophoretic mobility shift assay (EMSA)

EMSA detected the binding of fluorescently labeled Codanin1_Cterm_ and unlabeled CDIN1 with gradually increasing concentrations. Codanin1_Cterm_ (1.25 μM) fluorescently labeled by Alexa Fluor 488 was incubated with increasing amounts of CDIN1 (0-20 μM, 2-fold dilution line) in a final buffer (20 mM Tris-HCl, 150 mM NaCl, 5% glycerol, pH 8.0) for 15 min at room temperature. The binding reactions were resolved on native polyacrylamide gels (10% PAGE, 0.5×TBE). The runs were performed at 10 V cm^-1^ (Mini-PROTEAN spacer plate, Bio-Rad) for 45 minutes (or until loading dye reached the bottom of the gel) at 4 °C, and gels were visualized using a 473 nm laser and bandpass filter (510-530 nm) on a Typhoon™ FLA 9500 biomolecular imager (GE Healthcare Life Sciences). CDIN1-Codanin1_Cterm_ complex formation was observed as retardation in electrophoretic mobility of fluorescently labeled Codanin1_Cterm_.

### Size-exclusion chromatography (SEC)

Size-exclusion chromatography was used as a final refinement purification step to separate CDIN1 and Codanin1_Cterm_ from impurities of different sizes and to exchange buffers. Prior to size-exclusion chromatography, all protein samples were filtered through Nanosep filters CDIN1 (1 mg) and Codanin1_Cterm_ (0.75 mg) proteins were analyzed separately or as a complex preincubated for 15 minutes in a molar ratio 1:1, where the amount of the proteins was halved (0.5 mg and 0.375 mg) to achieve comparable signal levels. Chromatographic separation was performed on 10/300 GL Superdex 200 column (Cytiva) using a buffer (20 mM Tris-HCl, 150 mM NaCl, 5% glycerol, pH 8.0) as a mobile phase and flow rate 0.3 ml min^-1^. Fractions corresponding to the peaks from the chromatogram were collected and analyzed on 12% SDS-PAGE.

### Microscale thermophoresis (MST)

Microscale thermophoresis measured the binding affinity of a fluorescently labeled Codanin1_Cterm_ and CDIN1 by quantifying how CDIN1 affected the movement of a fluorescently labeled protein along a temperature gradient. The extent of binding was determined from the observed reduction in protein movement caused by the increase in the overall complex size upon binding. MST measurements were performed using a NanoTemper Monolith NT.115 (NanoTemper Technologies). Fluorescently labeled Codanin1_Cterm_ Alexa Fluor 488 (75 nM) was incubated for 15 min on ice with different concentrations of CDIN1 (0-5 μM, 2-fold serial dilutions), in a buffer (20 mM Tris-HCl, 150 mM NaCl, 5% glycerol, pH 8.0). The samples were loaded into premium-grade capillaries. MST assays were conducted at 25 °C, 100% blue-laser power, and medium MST power. The laser-on and laser-off intervals were 20 and 5 seconds, respectively. MO.Affinity Analysis v2.2.4 software was used to fit the data by Hill equation and to determine apparent K_D_ values from MST traces at 1.5 s time point after the infra-red laser irradiation. All measurements were performed at least three times for independently prepared sample sets.

### Isothermal titration calorimetry (ITC)

Isothermal titration calorimetry measured heat changes during the gradual titration of Codanin1_Cterm_ to CDIN1. The resulting thermogram provided the binding affinity, stoichiometry, and thermodynamic description of Codanin1_Cterm_ binding to CDIN1, without any protein labeling or modification. ITC experiments were conducted on a VPITC instrument (Microcal, GE Healthcare) at 15 °C. Proteins were dialyzed into the final buffer (20 mM TrisHCl, 150 mM NaCl, 5% glycerol, pH 8.0) and degassed before the experiment. The ITC cell (1 423 μl) was filled with CDIN1 (11 μM). Codanin1_Cterm_ (110 μM) was added in 20 consecutive titrations of 10 μl in 5-minute intervals, with a stirring rate of 240 rpm. Experimental data were analyzed in OriginPro 2022 version 9.9.0.225 software (OriginLab Corporation) using a one-site binding model to fit a titration curve. Binding enthalpy (ΔH), dissociation constant (K_D_), and reaction stoichiometry (n) were obtained from the fit.

### SEC Multiangle Light Scattering (MALS)

SEC-MALS combined the separation of molecules based on size through size exclusion chromatography (SEC), followed by measuring the intensity of scattered light at various angles (MALS) to provide information of size of the proteins and their complexes. CDIN1 (2 mg ml^-1^) and Codanin1_Cterm_ (2 mg ml^-1^) or complex CDIN1-Codanin1_Cterm_ in a ratio 1:1 were preincubated and dialyzed before the measurement in a buffer 20 mM Tris-HCl, 150 mM NaCl, pH 8.0 and then loaded onto an OmniSEC instrument (Malvern Panalytical) equipped with a 13-ml Zenix SEC 300 column (Sepax Technologies) previously equilibrated in the same buffer coupled to a Wyatt miniDAWN TREOS and Optilab T-rEX differential refractive index detector. Molar mass was calculated from the Raleigh ratio based on the multiangle static light scattering and protein concentration from the change in refractive index (dn/dc = 0.185). Analysis was performed using OMNISEC software 11.21 (Malvern Panalytical).

### Analytical ultracentrifugation (AUC)

Analytical ultracentrifugation at high orbital velocities monitored the sedimentation of CDIN1, Codanin1_Cterm_, and their complex in solution to determine molecular weight, shape, and native oligomeric states. AUC experiments were performed using ProteomeLab XL-I analytical ultracentrifuge (Beckman Coulter) equipped with an An-60 Ti rotor at 20 °C and 50 000 rpm. The sedimentation velocity (SV) experiment was performed in 12 mm titanium double-sector centerpiece cells (Nanolytics Instruments) loaded with 380 µl of both protein sample and buffer (20 mM Tris-HCl, 150 mM NaCl, pH 8.0). Scans were collected at 280 nm in 5-minute intervals at 0.003 cm spatial resolution in continuous scan mode. Protein samples were diluted in reference buffer to the final concentration of 24 μM CDIN1, 24 μM Codanin1_Cterm_, and complex CDIN1-Codanin1_Cterm_ in a ratio 1:1, where the concentration of the proteins was lowered to half (12 μM for both) to achieve similar signal levels. The partial specific volume of the protein and the solvent density and viscosity were calculated from the amino acid sequence and buffer composition, respectively, by the Sednterp software (http://bitcwiki.sr.unh.edu). The SV data corresponding to the complete sample sedimentation were fitted using the continuous c(s) distribution model in Sedfit v15.01c^13^. The M_W_ of particles was estimated based on the Svedberg equation, and c(s) distributions were normalized and plotted using the GUSSI software version 1 created by C. Brautigam (Univ of Texas Southwestern, Dallas, USA).

### SEC-Small-Angle X-ray Scattering (SAXS)

Size exclusion chromatography coupled with small angle X-ray scattering (SEC-SAXS) separated particles based on size and evaluated X-ray scattering of size-distributed fractions. SEC-SAXS revealed the size, shape, and low-resolution structure of CDIN1, Codanin1_Cterm_, and complex CDIN1-Codanin1_Cterm_ in solution. SEC–SAXS data of CDIN1-Codanin1_Cterm_ complex, CDIN1 and Codanin1_Cterm_ were collected at beamline P12 of the Petra III storage ring in the Deutsches Elektronensynchrotron (DESY, Hamburg, Germany) using Superdex 200 Increase 5/150 GL with flow 0.3 ml min^-1^ at an injection volume 45 µl and concentration of 10 mg ml^−1^ in 20 mM Tris-HCl, 150 mM NaCl, 5% glycerol, pH 8.0 buffer at 20 °C. Exposure time was 0.495 s per frame, and 1 200 frames were collected. Pilatus 6M detector was positioned at a sample-detector distance of 3 m and at a wavelength of λ = 0.123982 nm. All SAXS data were collected over the q range of 0.0259 – 4.434 nm^−1^.

The data were normalized to the intensity of the transmitted beam and radially averaged and subsequently processed with CHROMIXS, PRIMUS, AUTORG, and GNOM, as implemented in ATSAS v.3.2.1^14^. The frame sections for subsequent detailed analyses were selected using the Data Comparison tool within the ATSAS package and by examining the radius of gyration (Rg) distribution for CDIN1-Codanin1_Cterm_ complex. To identify oligomerically uniform frames, 10-frame segments of the SEC-SAXS peaks for Codanin1_Cterm_ and CDIN1 were analyzed by CRYSOL and OLIGOMER software^15^, respectively, using AlphaFold-predicted models for both the homodimer and monomer.

SAXS profiles were generated from selected intervals for CDIN1-Codanin1_Cterm_ complex (Frames 712 – 720), Codanin1_Cterm_ (Frames 842 – 851), and CDIN1 (Frames 692 – 701). The agreement between the SAXS data and AlphaFold-predicted CDIN1-Codanin1_Cterm_ heterodimer^16^, Codanin1_Cterm_ monomer, and CDIN1 homodimer were evaluated using CRYSOL^17^. The structures of CDIN1-Codanin1_Cterm_ heterodimer and CDIN1 homodimer were generated by ColabFold v1.5.3: AlphaFold2 using MMseqs2, replacing standard homology detection and MSA pairing. Experimental SAXS data of CDIN1, Codanin1_Cterm_, and complex CDIN1-Codanin1_Cterm_ and all related information compliant with SAXS data publication standards have been deposited in the Small Angle Scattering Biological DataBank (SASBDB)^18^ and Supplementary Table 1.

### Hydrogen deuterium exchange mass spectrometry (HDX-MS)

HDX-MS was used to monitor the exchange of backbone amide hydrogens with deuterium in CDIN1, Codanin1_Cterm_, and their complexes. Regions with reduced exchange were interpreted as protein interaction interfaces. CDIN1 and Codanin1_Cterm_ were diluted to a final concentration of 2 µM in H_2_O-based buffer (20 mM Tris-HCl, 150 mM NaCl, pH 8.0) to prepare non-deuterated controls and for peptide identification. The free individual CDIN1 and Codanin1_Cterm_, as well as complexed in the molar ratios of 1:2 and 2:1 (CDIN1 to Codanin1_Cterm_), were preincubated for 30 minutes at room temperature in H_2_O-based buffer and then diluted in D_2_O-based buffer (20 mM Tris-HCl, 150 mM NaCl, pH 7.6). To monitor HDX, samples were incubated at room temperature and quenched after 1 minute, 10 minutes, 30 minutes, or 1 hour, by adding 0.5 M TCEP-HCl, 4 M urea, 1 M glycine (pH 2.3) containing pepsin at a final concentration of 0.04 mg ml^-1^. Samples were then incubated for 3 min at 25 °C, rapidly frozen in liquid nitrogen, and stored at -80 °C before analysis. Final deuterium contents of the CDIN1 and Codanin1_Cterm_ buffer samples were 83% and 81%, respectively.

Frozen samples of CDIN1 and Codanin1_Cterm_ were thawed and injected at concentrations of 0.1 mg ml^-1^ and 0.07 mg ml^-1^, respectively, into a cooled LC-system (UltiMate 3000 RSLCnano, Thermo Scientific). Proteins were digested using two protease enzymatic columns (Nepenthesin-1 and pepsin, both 15 µl bed volume, Affipro s.r.o., CZ) connected in series. Initial pre-experimental measurements revealed insufficient sequence coverage with the dual column alone; therefore, an additional pepsin digestion step was introduced before freezing.

Peptides were trapped and desalted online on a peptide microtrap column (Michrom Bioresources, Auburn, CA) for 3 minutes. Both digestion and desalting were performed in a loading buffer (2% acetonitrile, 0.05% trifluoroacetic acid) at a flow rate of 100 µl min^-1^.

Next, the peptides were eluted onto an analytical column (Jupiter C18, 0.5 x 50 mm, 5 µm, 300 Å, Phenomenex, CA) and separated by a 29-minute linear gradient elution starting with 10% buffer B (80% acetonitrile in 0.08% formic acid) in buffer A (0.1% formic acid) and increasing to 40% buffer B at a flow rate of 50 µl min^-1^. The dual-protease, trap, and analytical columns were kept at 1.5 °C; initial trials at laboratory temperature indicated that storing the columns in a cooling box produced peptides of optimal lengths and significantly reduced back exchange.

Mass spectrometric analysis was carried out using an Orbitrap Elite mass spectrometer (Thermo Fisher Scientific) with ESI ionization connected online to a robotic system based on the HTS-XT platform (CTC Analytics, Zwingen, Switzerland). The instrument was operated in a data-dependent mode for peptide mapping (LC-MS/MS). Each MS scan was followed by MS/MS scans of the three most intensive ions from both collision-induced dissociation (CID) and higher-energy C-trap dissociation (HCD) fragmentation spectra. Tandem mass spectra were searched using SequestHT against the cRap protein database (ftp://ftp.thegpm.org/fasta/cRAP) containing the sequences of CDIN1 and Codanin1_Cterm_ recombinant proteins with the following search settings: mass tolerance for precursor ions of 10 ppm, mass tolerance for fragment ions of 0.6 Da, no enzyme specificity, two maximum missed cleavage sites and no-fixed or variable modifications. The false discovery rate at the peptide identification level was 1%. In the process of peptide mapping via LC-MS/MS, 242 peptides specific to CDIN1 and 232 peptides specific to Codanin1_Cterm_ were identified. For the final HDX-MS analysis, we used 108 peptides from each protein, CDIN1 and Codanin1_Cterm_. Sequence coverage was analyzed with Proteome Discoverer version 1.4 (Thermo Fisher Scientific) and graphically visualized with the MS Tools application^19^.

The measured raw files of all samples were evaluated using HDExaminer version 2.5 (Sierra Analytics, Modesto, CA). A peptide pool containing amino acid sequence, charge, retention time, and search-dependent score (Xcorr) was generated from the peptide-identified data. Deuterated and non-deuterated samples were then analyzed to calculate deuterium incorporation.

The software then evaluated the deuteration levels in protein regions and generated uptake plots showing cumulative differences in relative deuterium uptake (*ΔHDX = free protein - complex*). The uptake plots represented the deuteration of peptides over time with a calculated confidence level. Each accepted peptide was then mapped to the amino acid sequences of the analyzed proteins using the following procedure. Each residue was assigned uptake data only from peptides resolved with high confidence^20^. The ΔHDX value, expressed as a percentage (% ΔHDX), reflects the relative difference in HDX between the free and complexed state.

To cross-validate results, residue-level HDX values were also calculated using pyHDX software^21^. Residue deuteration levels were determined by weighted averaging of overlapping peptides, weighted by the inverse peptide length. ΔHDX values were used to identify potential interaction regions. The results for all time points were visualized as heatmaps with a calibrated color scale (Supplementary Figure 7).

To confirm system performance, HDX-MS analysis of a standard protein (myoglobin) was performed, as described previously^22^. Deuterium recovery was reproducible, and back-exchange levels were quantified following best practices^23^. The mass spectrometry proteomic data were deposited in the ProteomeXchange Consortium via PRIDE repository^24^ with the dataset identifier PXD037661.

### Co-immunoprecipitation (Co-IP)

Co-IP employed antibodies specifically recognizing Flag- and Myc-tagged CDIN1 and Codanin1 variants to capture a target protein, extracting other bound proteins to reveal protein-protein interactions. For Co-IP, HEK293T cells were cultivated at 37 °C and 5% CO_2_ using 12 ml of 1x Dulbecco’s Modified Eagle Medium per each 10-cm Petri dish – DMEM, high glucose, pyruvate supplemented, 4 mM L-glutamine, 1x MEM non-essential amino acids solution, 1x Penicillin-Streptomycin (all Gibco) and 10% FBS (Capricorn). When HEK293T cells reached almost complete confluency, cells were washed with 5 ml of 1x PBS and detached by the addition of 1 ml Trypsin-EDTA (0.25%), phenol red (both Gibco) for 1 minute, diluted in DMEM media, and split using one-third of the cells per 10-cm Petri dish. The next day, HEK293T cells were co-transfected with 10 μg plasmids containing Flag- and Myc-tagged protein constructs using polyethylenimine (PEI) on 10-cm Petri dishes.

The cells were harvested 48 hours after transfection, washed once with PBS, and lysed for 1.5 hours rotating at 4 °C in 500 μl of RIPA buffer (50 mM Tris-HCl, 150 mM NaCl, 0.5% Triton X-100, 1 mM EDTA, pH 7.4, with the addition of 1 mM DTT, 10 mM NaF, and protease inhibitor cocktail cOmplete tablets EDTA-free (Roche) right before use). After incubation, lysates were spun at 20 000 g for 15 min to remove debris. Supernatants were mixed with 40 μl of Anti-FLAG® M2 Magnetic Beads (Millipore) and incubated for 3 hours while rotating at 4 °C. The beads were washed three times in RIPA buffer without additives (50 mM Tris-HCl, 150 mM NaCl, 0.5% Triton X-100, 1 mM EDTA, pH 7.4) and three times with wash buffer (50 mM Tris-HCl, 150 mM NaCl, pH 7.4). For the subsequent elution, the beads were resuspended in 60 μl of elution buffer (50 mM Tris-HCl, 150 mM NaCl, pH 7.4) containing 3x Flag peptide in final concentration 150 μg ml^-1^ (Sigma-Aldrich) for 1.5 hours at 4 °C. Input supernatant and elution samples were separated on 12% SDS-PAGE and transferred onto a membrane (Cytiva, Amersham™ Protran®).

The following western blot analysis was done using Monoclonal ANTI-FLAG® M2-Peroxidase (HRP) antibody (Sigma-Aldrich), Monoclonal Anti-Myc tag antibody [9E10] (Abcam) produced in mice or Mouse Monoclonal Anti-alpha Tubulin antibody [DM1A] - Loading Control (Abcam). Flag antibody was diluted in a ratio of 1:3 000, Myc antibody was diluted in a ratio of 1:1 000 and Tubulin antibody was diluted in a ratio of 1:5 000 in blocking buffer (3.5% milk in TBS-T), and incubated O/N at 4 °C. Anti-Flag antibody was detected immediately after TBS-T wash (1x TBS, 0.1% Tween20). To detect anti-Myc and anti-Tubulin antibodies, the membrane was washed three times by TBS-T, incubated with secondary Anti-Mouse IgG (Fc specific) HRP (SigmaAldrich) diluted in a ratio of 1:5 000 for 1 hour at room temperature, and then washed three times again before detection by SuperSignal™ West Femto Maximum Sensitivity Substrate kit (Thermo Scientific) on Fusion FX instrument (Vilber).

### Circular dichroism (CD)

Circular dichroism determined the proportion of secondary structures in CDIN1 and Codanin1_Cterm_ by measuring the difference in the absorption of left and right circularly polarized light. Far-UV CD measurements were performed on a J-815 spectrometer (Jasco) at 20 °C in a 1-mm Quartz cuvette (Hellma Analytics). CD spectra of 0.2 mg ml^-1^ CDIN1 and 0.2 mg ml^-1^ Codanin1_Cterm_ proteins in a buffer (20 mM NaH_2_PO_4_, 150 mM NaF, pH 8.0) were acquired in the wavelength range of 190-250 nm with a 1 nm step at a scanning speed of 100 nm min^-1^. Each spectrum represents an average of ten accumulations. Subsequently, the buffer signal was subtracted, and data were converted from circular dichroism units to mean residue molar ellipticity (MRE) to account for precise protein concentration. The presence of secondary structural elements was evaluated using BeStSel software^25^ and compared with values obtained from AlphaFold models analyzed by PDBMD2CD software^26^.

### Differential scanning fluorimetry (nanoDSF)

Differential scanning fluorimetry determined the thermal stability – melting temperatures of CDIN1 and Codanin1_Cterm_ by monitoring fluorescence changes during heat-induced protein denaturation. NanoDSF measurements were carried out on Prometheus NT.48 instrument (Nanotemper Technologies) in nanoDSF standard grade capillaries. The measurements were performed in triplicates using 2 mg ml^-1^ CDIN1 and 1.5 mg ml^-1^ Codanin1_Cterm_ in a buffer (20 mM Tris-HCl, 150 mM NaCl, 5% glycerol, pH 8.0), the temperature in the range of 20-80 °C at a heating rate of 1 °C min^-1^ and excitation power 10%. Protein unfolding was monitored by fluorescence intensity measured at 330 and 350 nm. The melting temperature (T_m_) was determined from the first derivative of the fluorescence ratio (330/350)^27^.

### Dynamic light scattering (DLS)

Dynamic light scattering analyzed scattered light fluctuations caused by Brownian motion to determine the size of particles in solutions of CDIN1, and Codanin1_Cterm_. The homogeneity of protein samples was determined by DLS measurements using a Delsa Max Core (Beckman Coulter). Before the measurement, 4 mg ml^-1^ CDIN1 and 4 mg ml^-1^ Codanin1_Cterm_ in a buffer (20 mM Tris-HCl, 150 mM NaCl, 5% glycerol, pH 8.0) were centrifuged at 12 000 g, RT for 8 min. Twenty scans of 10 s data acquisition were averaged to examine the sample homogeneity. Data were collected and processed using software provided by Beckman Coulter. The homogeneity of samples and possible occurrence of aggregates was analyzed based on a fit of obtained autocorrelation functions of scattered light. Intensity-based data were qualitatively evaluated to determine aggregates indicated by peaks of *R*_h_ > 100 nm.

## Results

### Reconstituted CDIN1 and Codanin1Cterm are preferentially α-helical, homogenous and stable

We produced codon-optimized Codanin1_Cterm_ (amino acids 1005-1227) and full-length CDIN1 using a bacterial expression system. We purified both proteins using a three-step purification with a final step of size-exclusion chromatography (SEC). We assessed protein purity and verified their identity using SDS-PAGE and MALDI-TOF MS (Supplementary Figure 1).

To determine secondary structure composition, we combined circular dichroism measurements and predictions available in the AlphaFold Protein Structure Database^28^. We examined the predicted structures of CDIN1 (Q9Y2V0) and Codanin1 (Q8IWY9). The structural model for CDIN1 (Q9Y2V0) has been predicted with a high degree of certainty – out of the 281 amino acids, more than 257 showed a per-residue confidence score (pLDDT) above 90. CDIN1 predominantly forms α-helices with intermittent short β-strands, as shown in Fig. 1A. Moreover, the percentage of α-helix at 59% and β-sheet at 13% in the predicted model for CDIN1 aligns well with the circular dichroism (CD) measurements, which account for 45% α-helix and 13% β-sheet (Supplementary Figure 2A, B).

Likewise, the structural composition of Codanin1_Cterm_ has been reliably predicted, with a pLDDT score above 80 for 157 out of the 222 amino acids. According to the structural model, Codanin1_Cterm_ is arranged into α-helices that form two four-helix bundles (Fig. 1B). The predicted 53% α-helix content for Codanin1_Cterm_ closely correlates with the 55% helical structure content derived from our circular dichroism measurements (Supplementary Figure 2A, C). Altogether, structural predictions and our experimental CD data suggest that both CDIN1 and Codanin1_Cterm_ predominantly arrange into α-helices.

Additionally, to evaluate the size distribution of the purified proteins, we employed dynamic light scattering (DLS). The DLS plots (Supplementary Figure 2D) exhibited single peaks, suggesting that the proteins are monodisperse and free from aggregates. Finally, we used differential scanning fluorimetry (nanoDSF) to determine the thermal stability of CDIN1 (48 °C) and Codanin1_Cterm_ (55 °C) (Supplementary Figure 2E), suggesting that both proteins are stable at room temperature. Taken together, we prepared pure proteins that were adequately folded preferentially into α-helices, homogenous, stable, and thus well-suitable for subsequent studies.

### CDIN1 and Codanin1_Cterm_ form complex with 1:1 stoichiometry

After successfully purifying and characterizing CDIN1 and Codanin1_Cterm_, we examined their complex formation qualitatively. To reveal CDIN1-Codanin1_Cterm_ interacting stoichiometry and complex composition, we used an electrophoretic mobility shift assay (EMSA) (Fig. 1C). In the EMSA experiment, we incrementally added CDIN1 to the fluorescent-tagged Codanin1_Cterm_. Notably, a strong signal consistent with a heterodimer was detected in all lanes where CDIN1 concentration exceeded 300 nM. Accordingly, a band corresponding to free Codanin1_Cterm_ vanished upon achieving a 1:1 molar ratio of CDIN1:Codanin1_Cterm_, indicating an equal stoichiometry of the complex. Additionally, we detected a slowly migrating fraction of molecules arranged into a possible heterotetrameric complex (Fig. 1C).

When examining the complex using size-exclusion chromatography (SEC), we also identified two distinct peaks, suggesting that CDIN1 and Codanin1_Cterm_ form heterodimeric and heterotetrameric complexes (Fig. 1D). Moreover, semi-quantitative SDS-PAGE densitometry analysis of SEC fractions corresponding to peak maxima revealed the comparable intensities of CDIN1 and Codanin1_Cterm_ bands (see lanes 1 and 2 in the inset of Fig. 1D). The similar intensities of both proteins support the equimolar content of CDIN1 and Codanin1_Cterm_ in the complex.

Simultaneously, we quantified the molecular weight of complexes CDIN1 and Codanin1_Cterm_ separated by size and shape using SEC coupled with multiangle light scattering (SEC-MALS). The most pronounced SEC peak aligns with a molecular weight of 60 kDa (Fig. 1E), matching the anticipated molecular weight of 58 kDa for the heterodimer of CDIN1 and Codanin1_Cterm_. The broader, less pronounced SEC peak contains complex fractions with molecular weights in the range 112-130 kDa that well accommodate heterotetramer comprising two CDIN1 and two Codanin1_Cterm_ molecules with an anticipated total molecular weight of 118 kDa.

Next, we used analytical ultracentrifugation (AUC) to quantify how proteins assemble into complexes. The sedimentation AUC profiles (Fig. 1F) corroborate that CDIN1 and Codanin1_Cterm_ preferentially form heterodimeric and potentially tetrameric complexes. In summary, the sedimentation observations agree with SEC-MALS results, showing the preference for heterodimer CDIN1:Codanin1_Cterm_ formation in 1:1 ratio.

### CDIN1 binds Codanin1_Cterm_ tightly with nanomolar affinity

To quantify the interaction between CDIN1 and Codanin1_Cterm_, we carried out microscale thermophoresis (MST) and isothermal titration calorimetry (ITC) measurements. During MST analysis (Fig. 1G), we observed a change in fluorescence intensity of labeled Codanin1_Cterm_ upon the addition of CDIN1. The MST single-inflection sigmoidal curve fit revealed that CDIN1 binds Codanin1_Cterm_ with a K_D_ value of 53 ± 8 nM.

To validate the binding affinity, we performed calorimetric titration with the complementary arrangement (Fig. 1H) – Codanin1_Cterm_ was gradually injected into the calorimetric cell containing CDIN1. ITC thermograms provided a K_D_ value of 20 ± 4 nM for Codanin1_Cterm_ binding to CDIN1. The partial oligomerization of individual proteins could explain why the stoichiometry obtained from ITC measurements is below the expected equimolar ratio and the difference in the obtained affinity values determined by ITC and MST. Overall, we can confidently state that CDIN1 and Codanin1_Cterm_ exhibit high affinity with K_D_ value notably below 100 nM, as verified by two independent quantitative methods.

### Predicted heterodimeric complex, primarily monomeric Codanin1_Cterm_, and dimeric CDIN1 align with experimental SEC-SAXS and SEC-MALS profiles

To further experimentally examine how CDIN1 and Codanin1_Cterm_ are assembled into a complex and arranged in solution, we carried out SEC coupled with small-angle X-ray scattering (SAXS). We separated discrete oligomeric states by SEC and compared their SAXS profiles with predicted models (Fig. 2). The Guinier plot for Codanin1_Cterm_ demonstrates linearity (Supplementary Figure 4C), indicating no aggregation or inter-particle interference in the solution. Conversely, there is evidence of radiation-induced aggregation in the initial parts of Guinier plots of the complex (Supplementary Figure 3C) and CDIN1 (Supplementary Figure 5C), suggesting lower stability under beam exposure. Nonetheless, the observed radiation damage did not impede further analysis of the SAXS data (Supplementary Figure 3-5).

**Fig. 2:**
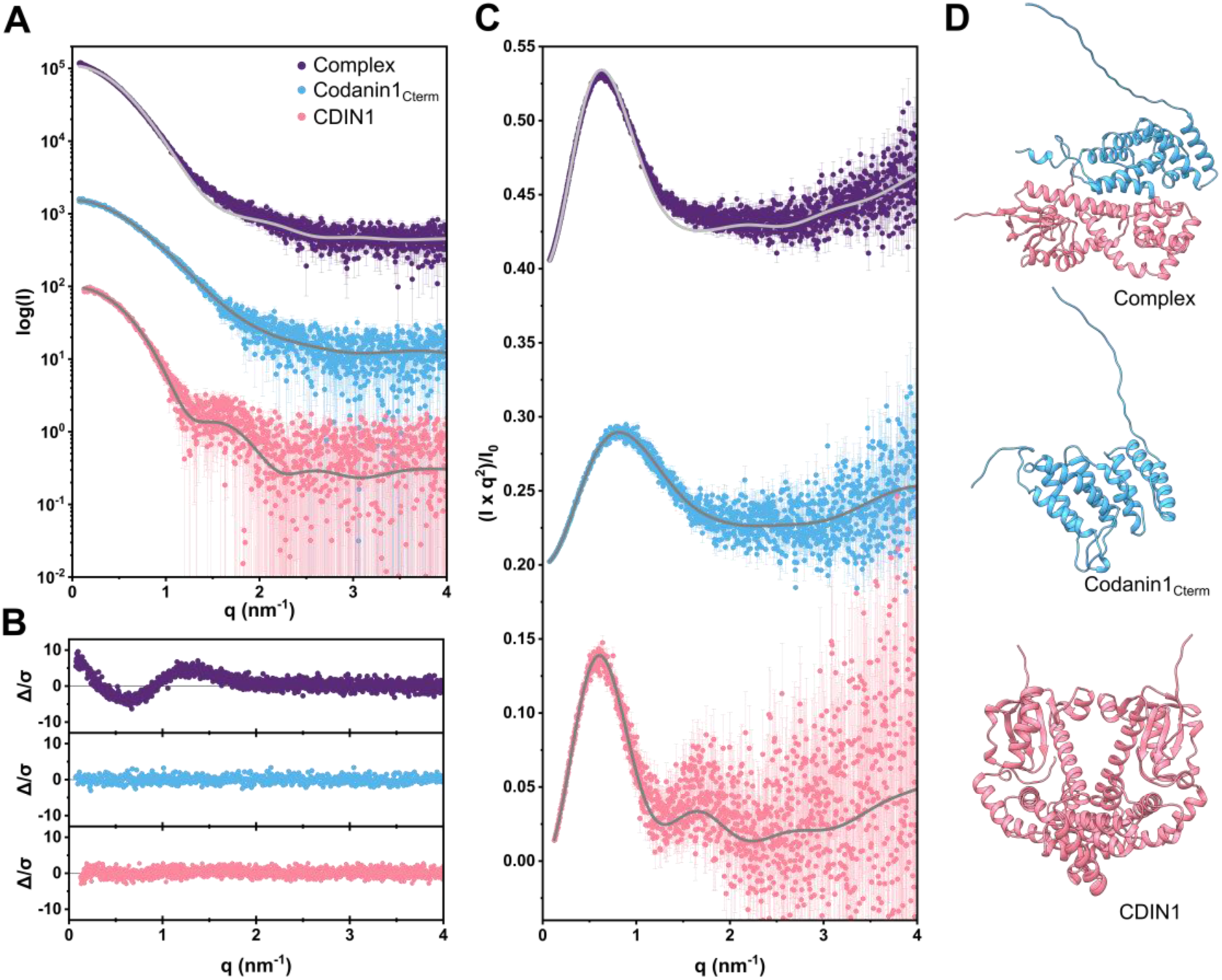
SEC-SAXS data along with predicted structures of Codanin1_Cterm_, CDIN1, and their complex. (A) SAXS profiles of CDIN1-Codanin1_Cterm_ complex (purple), Codanin1_Cterm_ (blue), and CDIN1 (pink) (I is scattering intensity; q is momentum transfer) overlaid with the profiles (gray) calculated by CRYSOL, from the predicted AlphaFold models for CDIN1-Codanin1_Cterm_ complex, Codanin1_Cterm_ and CDIN1. I(q) versus q plots are offset vertically. (B) Error-weighted residual plots for the CRYSOL model fits of CDIN1-Codanin1_Cterm_ complex, Codanin1_Cterm_, and CDIN1. (C) Dimensionless Kratky plots show the globularity of studied proteins. The plots are stacked vertically, so each panel is shifted by 0.2. (D) AlphaFold predicted models of heterodimeric CDIN1-Codanin1_Cterm_ complex, monomeric Codanin1_Cterm_, and homodimeric CDIN1 used by CRYSOL to fit the experimental SAXS data. Error bars are standard errors that might be smaller than the symbols and therefore covered.

Simultaneously, we quantified the distribution of CDIN1 and Codanin1_Cterm_ by combining size exclusion chromatography and multiangle light scattering (SEC-MALS). For CDIN1- Codanin1_Cterm_ complex, we observed a prominent SEC peak within the frame range 680 – 900 (Supplementary Figure 3). Using the Data Comparison tool within ATSAS package, alongside a thorough examination of the distribution of the radius of gyration (R_g_), we selected frame numbers 712 – 720 with R_g_ plateau for detailed SAXS analysis (highlighted in Supplementary Figure 3A). We determined R_g_ using the Guinier approximation and by analyzing the concentration dependence of the distance distribution function P(r), with both methods yielding consistent R_g_ values of 3.04 ± 0.01 nm. The observed molecular weight of 57 kDa matches the anticipated 58 kDa (Supplementary Figure 1), indicating CDIN1-Codanin1_Cterm_ heterodimer formation. Additionally, the Kratky plot exhibits a bell-shaped curve, indicating the formation of folded globular protein structures. We used CRYSOL to match AlphaFold predicted models with the experimental SAXS profile of CDIN1-Codanin1_Cterm_ complex (Fig. 2). The obtained Chi-Square (Χ^2^) value of 6.1 implies the difference in predicted structure and experimental SAXS data that might reflect the free spatial movement of unstructured terminal parts of Codanin1_Cterm_ along with flexible C-terminal part of CDIN1. The partial complex sample heterogeneity shown by SEC-SAXS was also evident in previous SEC-MALS and AUC experiments (Fig. 1E, F).

For Codanin1_Cterm_, the major peak eluting from gel filtration column corresponded to the Codanin1_Cterm_ monomer as judged by molecular weight estimates from SAXS – 30 kDa (Supplementary Table 1) and MALS – 26 kDa (Supplementary Figure 2F), close to the anticipated value of 25 kDa (Supplementary Figure 1D). Additionally, the SAXS profile of Codanin1_Cterm_ gave linear Guinier region and bell-shaped P(r) functions (Supplementary Figure 4C, D) with the R_g_ and D_max_ values aligned with a monomer structure predicted by AlphaFold (Fig. 2). Comparison of the experimental scattering curve and scattering pattern calculated from the Codanin1_Cterm_ monomeric model provided X^2^ = 1.0 and for the homodimer of Codanin1_Cterm_ X^2^ = 10.4 (Supplementary Figure 4E, F) further indicated that Codanin1_Cterm_ adopts preferentially a monomeric state.

When we analyzed CDIN1 by SEC-SAXS, we detected a single peak, exhibiting variant R_g_ values throughout the peak suggesting the presence of various oligomeric states of CDIN1 (Supplementary Figure 5). We examined highlighted 10-frame segments of the peak using OLIGOMER software along with AlphaFold-predicted models for both the dimer and monomer of CDIN1 (Supplementary Figure 5A). The evaluation of the early eluting part of the peak with consistent R_g_ values, within frames 692 - 701 (Supplementary Figure 5), revealed a composition 100% of dimers and 0% monomers. Consequently, we compared the experimental scattering curve and calculated scattering pattern from the CDIN1 homodimeric model, yeilding X^2^ = 1.06 (Fig. 2). Conversely, the evaluation of the descending limb peak segment in frames 841 - 851 indicated a composition of 32% dimers and 68% monomers; Supplementary Figure 5A includes an inset plot showing the distribution of the dimeric state of CDIN1 across the frames of SEC-SAXS peak. The Kratky plot shows that CDIN1 dimer has a globular overall shape; however, the presence of a minor peak at around the scattering factor of 1.75 nm^-1^ (Fig. 2 A, C) suggests the occurrence of two globular subdomains, which are also noticeable as two lobes in the AlphaFold model of CDIN1 dimer (Fig. 2 D).

The MALS analysis of the most pronounced SEC peak of CDIN1 revealed a molecular weight of 69 kDa (Supplementary Figure 2G); SAXS data provided 66 kDa (Supplementary Table 1), matching the anticipated weight of a homodimer – 67 kDa (Supplementary Figure 1C). The broad peak with a maximum corresponding to 49 kDa might reflect the equilibrium between the monomeric and dimeric state of CDIN1. We also detected a signal at 137 kDa that might correspond to a homotetrameric arrangement of CDIN1.

In conclusion, our SEC-MALS and SEC-SAXS measurements suggest that CDIN1 adopts predominantly homodimeric arrangements in solution, whereas Codanin1_Cterm_ occurs primarily as a monomer, and together CDIN1 and Codanin1_Cterm_ form heterodimers.

### CDA-I-related mutations residing in interacting regions disturb CDIN1-Codanin1_Cterm_ binding

To identify interacting regions of CDIN1 and Codanin1_Cterm_, we carried out hydrogen deuterium exchange (HDX) coupled with mass spectrometry (MS). Based on tandem mass spectrometry detection connected with liquid chromatography (LC-MS/MS) bottom-up proteomics analysis of both proteins, the peptides covering whole protein sequences were identified. To evaluate HDX-MS data, we created the library from the peptides with high confidence, which covered 100% of Codanin1_Cterm_ and 89% of CDIN1 (Supplementary Figure 6). We prepared samples of each protein alone or saturated with a two-fold excess of the interacting partners. We quantified the difference in relative deuterium uptake (ΔHDX) by comparing the exchange behavior of free individual proteins and complexed proteins samples in the same D₂O-based buffer at four time points (1 minute, 10 minutes, 30 minutes, and 1 hour)^20,22^. The 10-minute time interval was chosen as a representative point for illustrating the HDX-MS profiles (Fig. 3B, E). The resulting differential deuterium uptake plots indicated interacting regions that were less exposed to solvent upon binding. To interpret the data comprehensively, we used modified Woods plots displaying the length and positions of residues of each analyzed peptide against their respective difference in relative deuterium uptake^29^. Deuterium uptake heatmaps for all incubation times are shown in Supplementary Figure 7 and the PRIDE repository^24^.

**Fig. 3.**
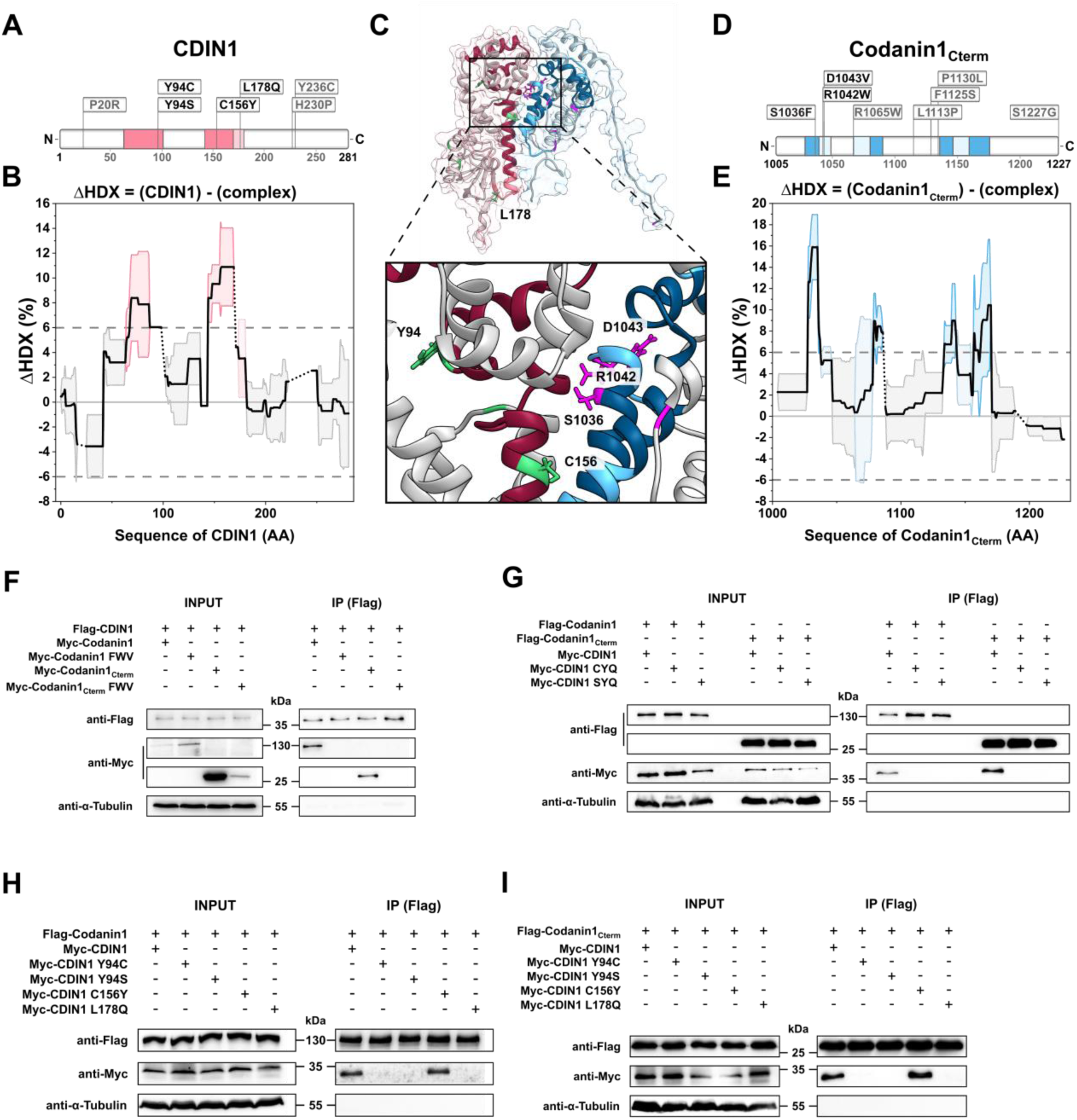
CDA-I-associated mutations in interacting regions disrupt CDIN1 and Codanin1 binding. (A, D) Scheme of CDIN1 and Codanin1_Cterm_ highlighting CDA-I-associated mutations and interacting regions revealed by HDX-MS. (B, E) Significant sum differences in relative deuterium uptake (*ΔHDX = CDIN1 - complex*) for CDIN1 upon the addition of two-fold molar excess of Codanin1_Cterm_, and differences in relative deuterium uptake (*ΔHDX = Codanin1_Cterm_ - complex*) for Codanin1_Cterm_ upon the addition of two-fold molar excess of CDIN1 after 10 minutes of deuteration, respectively. Highlighted colored regions represent interacting sites with mean value (dark colored) or standard deviation (light colored) exceeding the 6% deuteration threshold, dotted areas represent regions with incomplete peptide coverage. (C) Heterodimer CDIN1-Codanin1_Cterm_ structure predicted by AlphaFold. Interacting regions determined by HDX-MS are colored-coded. Dark shading indicates regions with mean value, light shading denotes regions with standard deviation exceeding the 6% deuteration threshold – red for CDIN1 and blue for Codanin1_Cterm_; noninteracting regions are shown in gray; CDA-I-associated mutations are marked in green for CDIN1, in magenta for Codanin1_Cterm_. The mutated amino acids in interacting regions are displayed as sticks and numbered. (F, G) Complementary Co-IP analyses using wild type, triple-mutated Codanin1 and CDIN1 proteins expressed in HEK293T cells show that simultaneous introduction of CDA-I-associated mutations in either Codanin1 or CDIN1 interacting regions disrupts CDIN1-Codanin1 binding. (H, I) Co-IP analyses using proteins expressed in HEK293T cells identify individual CDIN1 mutations associated with CDA-I that are critical for disrupting the binding to full-length Codanin1 or Codanin1_Cterm_.

We identified two regions in CDIN1 that significantly prevented deuterium uptake upon binding (elevated ΔHDX): amino acids 63-98, and 143-172 (Fig. 3A, B). In Codanin1_Cterm_, we identified three regions with reduced deuteration (less solvent-exposed region), suggesting interaction sites of CDIN1 and Codanin1_Cterm_ complex: amino acids 1028-1046, 1064-1087, and 1133-1170 (Fig. 3D, E).

Interestingly, we found that the interacting regions identified by HDX-MS contain mutations that have been described in CDA-I patients (Fig. 3A, D). Specifically, in interacting regions of CDIN1, we localized CDA-I-related mutations Y94C, Y94S, C156Y, and L178Q. In Codanin1_Cterm_ interacting sequence, we found three disease-associated mutations S1036F, R1042W, and D1043V.

To evaluate how CDA-I-related mutations residing in interacting regions affect the mutual binding of CDIN1 and full-length Codanin1, we employed site-directed mutagenesis followed by co-immunoprecipitation (Co-IP). We examined the interaction of CDIN1 and Codanin1 using proteins expressed in HEK293T to ensure that we analyzed full-length proteins with post-translational modifications as they occur in human cells. Crucially, our Co-IP results revealed that both full-length Codanin1 and truncated Codanin1_Cterm_ bind CDIN1, providing strong support for using truncated Codanin1_Cterm_ in experiments describing interaction with CDIN1 (Fig. 3F, G). We carried out additional control experiments to confirm the specificity of the used antibodies (Supplementary Figure 8).

When we introduced a cluster of three closely located mutations S1036F-R1042W-D1043V (FWV) into Codanin1 or Codanin1_Cterm_ (Fig. 3C), we observed no detectable interaction with CDIN1 (Fig. 3F), suggesting that the mutation cluster in Codanin1 prevents its binding to CDIN1. In the complementary experimental setup, we investigated how mutations in interacting regions of CDIN1 affect binding to Codanin1. Initially, we introduced two triplets of mutations Y94C-C156Y-L178Q (CYQ) and Y94S-C156Y-L178Q (SYQ) into CDIN1 (Fig. 3C). We found that introducing mutation triplets into CDIN1 aborted binding to Codanin1 (Fig. 3G).

Furthermore, to identify which mutations are critical for CDIN1 binding to Codanin1, we prepared CDIN1 variants with single mutations. Our Co-IP experiments revealed that exchanging hydrophobic tyrosine to polar cysteine or serine at CDIN1 position 94 in mutants Y94C and Y94S disrupted binding to Codanin1. Similarly, exchanging hydrophobic leucine for negatively charged glutamine in the L178Q mutant of CDIN1 interrupted binding to Codanin1. On the contrary, introducing tyrosine instead of cysteine in C156Y mutant of CDIN1 had no effect on Codanin1 binding (Fig. 3H, I). Thus, three of four mutations in the interacting region of CDIN1 are individually sufficient to abolish binding with Codanin1. In summary, we demonstrated that CDA-I-related mutations in interacting regions impair binding of CDIN1 and Codanin1.

## Discussion

Here, we discuss the structural and functional implications of the presented findings of molecular arrangements of CDIN1 and Codanin1_Cterm_ that bind tightly and the finding that the interacting regions of both proteins reside CDA-I-associated mutations that disrupt CDIN1 and Codanin1_Cterm_ interaction. We also present future research directions based on our findings.

### Implications for molecular arrangements and interactions

Our combined structural studies of individual proteins and the complex have indicated that CDIN1 and Codanin1_Cterm_ bind to form predominantly a heterodimer, while individual CDIN1 molecules preferentially form dimers. The prevalent dimeric state of CDIN1 has been detected by three different methods in a wide range of concentrations – AUC (0.6 mg ml^-1^), SEC-MALS (2 mg ml^-1^), and SEC-SAXS (injected at 10 mg ml^-1^). However, the SAXS profiles that agree well with AlphaFold predicted models suggest that CDIN1 does not retain a homodimeric form when interacting with Codanin1_Cterm_ in the heterodimeric complex (Fig. 2D).

The observed differences between the SAXS data and the AlphaFold model of CDIN1-Codanin1_Cterm_ complex (Fig. 2, Supplementary Figure 3) may arise from several factors. Firstly, the model itself may not fully capture the true structure of the complex, potentially due to limitations in the predictive accuracy of AlphaFold for this specific system. Secondly, the presence of multiple species in solution, such as monomers, dimers and higher order multimers, could contribute to the observed differences. The complex heterogeneity in the solution state can lead to an average SAXS signal that does not correspond to a single, well-defined structure. Lastly, conformational heterogeneity within the proteins, including flexible or disordered regions, may result in a range of conformations that are not adequately represented by a single static model. These factors collectively indicate that the SAXS data for CDIN1-Codanin1_Cterm_ complex represents a more intricate and dynamic array of structures than what the current AlphaFold model can depict and should be interpreted with appropriate caution. Despite the limitations discussed, the evidence from complementary techniques consistently supports the prevalence of the heterodimeric state of the complex, reinforcing the validity of this finding.

On the contrary, we showed with high experimental confidence that Codanin1_Cterm_ occurs mainly as a monomer (Fig. 2, Supplementary Figure 4). We should nevertheless consider the limitations of our findings for Codanin1_Cterm_, as we analyzed the separate C-terminal region of Codanin1 that might be affected by the other regions of full-length Codanin1.

We demonstrated that CDIN1 and Codanin1_Cterm_ bind directly in vitro. The direct binding of CDIN1 and Codanin1 is consistent with previous immunoprecipitation studies^2,10^. The determined binding affinity of CDIN1 and Codanin1 is comparable with other chromatin-shaping proteins^30–32^. We found that CDIN1 and Codanin1_Cterm_ form complexes in equimolar ratio based on stoichiometry determined from EMSA experiments (Fig. 1C), the densitometry analysis of PAGE profiles corresponding to SEC peaks (Fig. 1D, inset), and ITC titration (Fig. 1H).

### Functional implications of CDA-I-related mutations

After successfully defining interacting regions by HDX-MS, we revealed by co-immunoprecipitation that CDA-I-associated mutations in interacting regions of CDIN1 and Codanin1 disrupt their mutual binding (Fig. 3). We identified three disease-related mutations in CDIN1 that individually diminish CDIN1’s ability to bind to Codanin1. Furthermore, we identified a mutation triplet in Codanin1 that prevents CDIN1 binding. The mutations in both proteins affect the binding between CDIN1 and Codanin1, but the interaction network is much wider, when we examine how Codanin1 connects with ASF1.

Recent studies elucidated the atomic structure of the ASF1-Codanin1 complex using cryo-electron microscopy (cryo-EM) ^33,34^. Interestingly, the cryo-EM structures reveal that Codanin1 dimer facilitates two ASF1 subunits via a dual-binding mode involving previously identified B-domain and newly described Histone-mimicking helix (HMH). B domain and HMH are highly sequentially conserved across species, highlighting their evolutionary and functional significance in ASF1 regulation by Codanin1^34^. Codanin1 can engage both ASF1 interaction sites simultaneously. First, Codanin1 competes with other histone chaperones, CAF-1 and HIRA, for the binding site on ASF1 using B-domain. Second, Codanin1 HMH motive occupying the histone binding site of ASF1 implies that Codanin1 prevents ASF1 from binding H3-H4 histone dimers. Thus, Codanin1 binding ASF1 through HMH motive would act as a potent inhibitor of ASF1-mediated histone supply.

Structural data also showed that the C-terminal region of Codanin1 is flexible. Biochemical analysis together with structural prediction, revealed that the C-terminal region is also involved in Codanin1 dimerization. Additionally, the C-terminal domain of Codanin1 is necessary for interaction with CDIN1^34^. Nevertheless, the structure of the Codanin1’s C-terminal part that interacts with CDIN1 remains enigmatic due to high flexibility. To address the structural flexibility, the authors used AlphaFold predictions, which resulted in a model that closely aligns to the model validated by our SAXS measurements.

Even though, the CDA-I-related mutations are directly connected with patients suffering from CDA-I disease, the molecular effect of individual mutations remains elusive. The suggestion of the molecular effect is suggested by the authors where they showed that pathogenic CDA-I mutation in Codanin1 (P672L) affect ASF1 sequestration^34^. Moreover, CDA-I-associated mutations, such as R714W and R1042W, might compromise the interaction between Codanin1 and ASF1 ^7^.

Our results raise the fundamental question: what is the molecular function of CDIN1? We speculate that CDIN1, via its binding to the C-terminal part of Codanin1, may stabilize Codanin1.

To evaluate the contribution of structured regions, we aligned the AlphaFold predicted CDIN1 and Codanin1_Cterm_ secondary structures (Fig. 1A, B) with our HDX-MS data (Fig. 3 A, D). The regions with the most reduced deuteration in the protein complex, compared to the free form of the proteins, indicate interaction sites confirmed by co-immunoprecipitation (Supplementary Figure 7). However, our measurements do not exclude the possible role of the disordered regions in full-length Codanin1. For example, CDIN1 could potentially stabilize full-length Codanin1 by rearranging the disordered regions into a more condensed structure, thereby augmenting the structural stability of Codanin1.

Additionally, we hypothesize that the CDIN1-mediated structural arrangement of Codanin1 might be critical for the effective regulation of undisturbed histone trafficking, which is essential for proper chromatin formation. The disturbance of tightly controlled histone depositions through ASF1 sequestration by Codanin1 might contribute to the occurrence of spongy “Swiss cheese“ chromatin and internuclear bridges in erythroblasts – the typical hallmarks of CDA-I disease (Fig. 4). We present a theoretical model explaining the role of CDIN1-Codanin1 complex in erythroblast development and its implication for CDA-I. Our hypothetical model is in accordance with the regulatory mechanism suggested by the Groth laboratory^7,34^, where Codanin1 binding is limiting step in ASF1-mediated nuclear import of histones H3/H4.

**Fig. 4:**
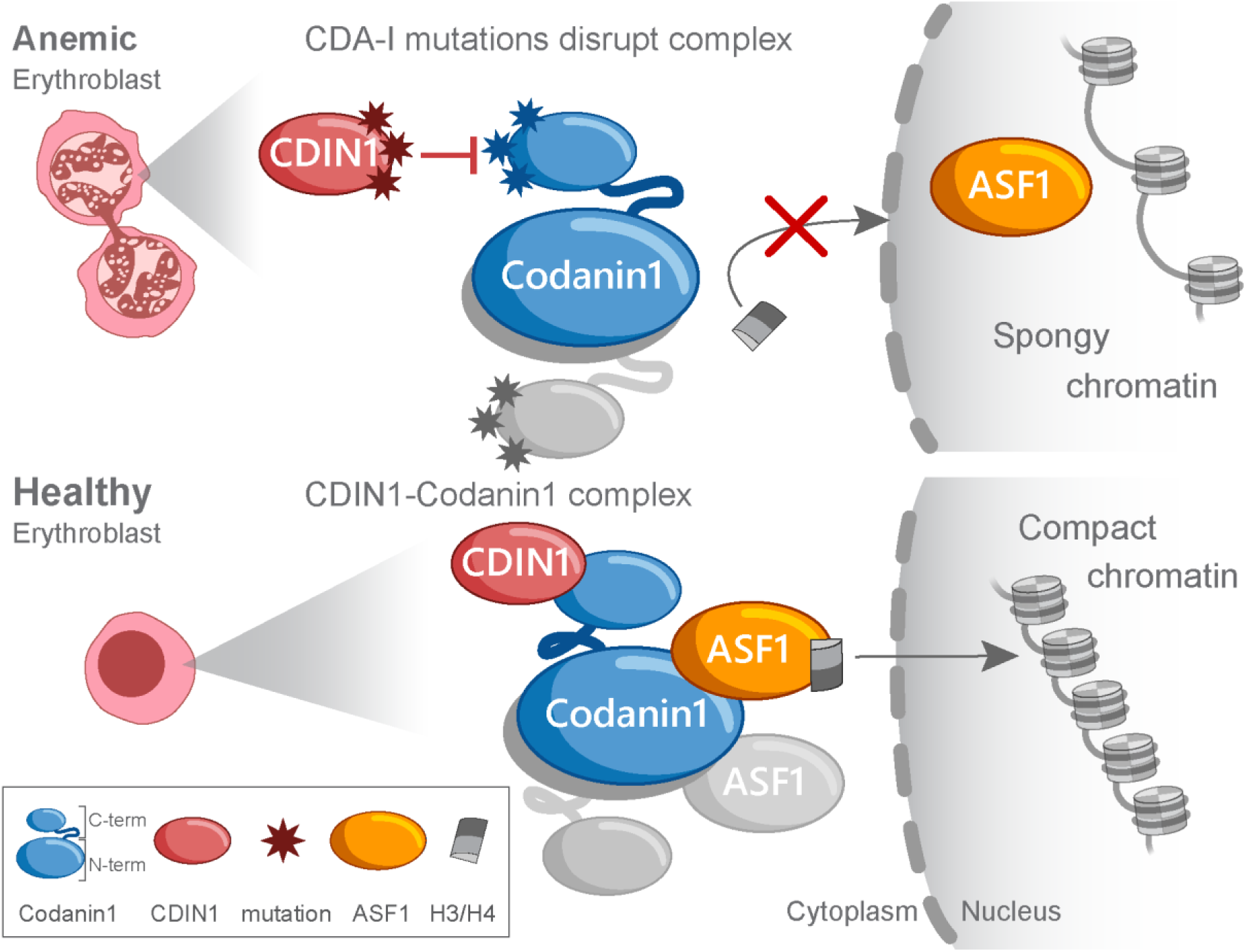
The hypothetical role of CDIN1 and Codanin1 in CDA-I progression and heterochromatin compaction. A putative model of how CDIN1-Codanin1 complex affects the arrangement of chromatin in erythroblasts suggests that if CDA-I-associated mutations prevent CDIN1-Codanin1 complex formation thus prevent ASF1 sequestration to cytoplasm. Hence, ASF1 is unable to traffic H3-H4 histones from cytoplasm to nucleus. Lack of histones in the nucleus leads to a loosely arranged anemic heterochromatin, which compromises erythropoiesis, as abnormal erythroblasts with internuclear bridges indicate. Conversely, wild type CDIN1 binds Codanin1, which has no effect on ASF1’s transport of H3/H4 histones to the nucleus, resulting in proper heterochromatin compaction, allowing normal erythropoiesis and the formation of healthy erythroblasts.

We further expand the molecular model of CDA-I progression by proposing that Codanin1 through its B-domain and HMH domain binds ASF1 in the cytoplasm. By structural rearrangements ASF1 facilitates H3-H4 dimer binding and the Codanin1-ASF1-H3-H4 complex is subsequently transferred to nucleus. The well-compacted chromatin leads to normal erythroblast maturation and function. However, when CDA-I mutations impair CDIN1 binding to Codanin1, Codanin1 is unable to sequester ASF1 to cytoplasm for binding and transporting H3-H4 dimer to the nucleus. The histone deficiency in the nucleus leads to a defective heterochromatin structure and impaired erythropoiesis, as manifested by the presence of abnormal erythroblasts in CDA-I patients.

### Future research directions

To elucidate whether and how CDIN1 affects the stability of Codanin1, further investigations involving full-length Codanin1 are warranted. Our interaction studies indicate a preferential binding of CDIN1 to the C-terminal part of Codanin1, even in the presence of other Codanin1 domains.

Furthermore, we used AlphaFold 3^35^ to predict the interaction of ASF1a and the full-length Codanin1. In Supplementary Figure 9, we present the alignment of the predicted ASF1a-Codanin1 complex with previously published crystallography data of ASF1-HIRA^36^ and ASF1-Histone H3/H4^37^, which is in complete agreement with recently published Cryo-EM structures^33,34^. The revealed structural alignment shows that Codanin1 wraps tightly around ASF1a. Thus, Codanin1 could prevent ASF1 from binding to HIRA and Histones H3/H4. The ability of Codanin1 to interact with both CDIN1 and ASF1 motivates further investigation of how CDIN1 and ASF1 affect the function of full-length Codanin1 that even recent cryo-EM data has not addressed yet^33,34^.

Finally, follow-up research utilizing erythroblast model cell lines must be conducted to determine if specific mutations that disrupt CDIN1-Codanin1 binding also cause a phenotype typical of CDA-I. The proposed research directions will provide additional deeper insights into the underlying mechanisms of the molecular origin of inherited anemia CDA-I.

In summary, our research offers a framework for the structural understanding of how CDIN1 and Codanin1 interact and thus contribute to chromatin arrangement that must be tightly handled during DNA replication and gene transcription. Here we quantified the interaction of CDIN1 and Codanin1 and defined interacting regions. We identified CDA-I-related mutations in CDIN1 and Codanin1, disturbing the mutual interaction of proteins tightly associated with CDA-I progression. Our findings of structural arrangements of CDIN1 and Codanin1 are an important step towards elucidating the molecular origin of CDA-I disease.

## Data availability

The small-angle X-ray scattering data have been deposited to the Small-angle scattering biological data bank (SASBDB)^18^: CDIN1-Codanin1_Cterm_ complex – <u>SASDVJ4</u>, Codanin1_Cterm_ – <u>SASDV49</u>, and CDIN1 – <u>SASDV39</u>. The hydrogen deuterium exchange mass spectrometry proteomics data have been deposited to the ProteomeXchange Consortium via the PRIDE repository^24^ with the dataset identifier PXD037661. Supplementary Data are available online.

## Acknowledgments

The Czech Science Foundation (GA23-05241S to C.H.) has primarily supported this research. The research has been carried out with institutional support of the Institute of Biophysics of the Czech Academy of Sciences (68081707 to M.S., T.B., and P.V.). This work benefited from access to the EMBL Hamburg, an Instruct-ERIC centre. Financial support was provided by Instruct-ERIC (PID 27193), CIISB, Instruct-CZ Centre of Instruct-ERIC EU consortium, funded by MEYS CR infrastructure project LM2023042 and European Regional Development Fund-Project “UP CIISB” No. CZ.02.1.01/0.0/0.0/18_046/0015974 is gratefully acknowledged for the financial support of the measurements at the following core facilities: CEITEC – Proteomics, Biomolecular Interactions and Crystallography, CMS-Biocev – Biophysical techniques, Crystallization, Diffraction, and Structural mass spectrometry. The research has been supported by the National Institute for Cancer Research (Programme EXCELES, ID Project No. LX22NPO5102) – Funded by the European Union – Next Generation EU and by the Ministry of Health, Czech Republic – Conceptual Development of Research Organization, MH CZ - DRO (MMCI, 00209805). T.B. is supported by Brno Ph.D. Talent Scholarship – funded by Brno City Municipality.

We recognize Agnel Sfeir for her brilliant, inspiring ideas and wholehearted support on both scientific and personal levels. We thank Samuel Tremblay-Belzile for his critical reading and editing of the manuscript. We thank Melissa Graewert for her insightful comments and advice on SAXS data evaluation.

## Author contributions

M.S. designed the initial studies, performed the biophysical characterization and immunoprecipitation studies, and wrote the manuscript. T.B. designed and carried out SEC-SAXS, SEC-MALS, and AUC experiments and wrote the manuscript. I.N. prepared and characterized initial protein constructs, T.J. designed initial interaction experiments, P.V. cloned and designed protein expression constructs, L.H., N.V. and L.U. carried out HDX-MS measurements and analyzed interaction mapping data, C.H. designed the study and wrote the manuscript. All authors read and approved the manuscript.

## References

1. Iolascon, A., Andolfo, I. & Russo, R. Congenital dyserythropoietic anemias. Blood 136, 1274–1283 (2020).

2. Olijnik, A.A. et al. Genetic and functional insights into CDA-I prevalence and pathogenesis. Journal of Medical Genetics 58, 185–195 (2021).

3. Wickramasinghe, S.N. & Wood, W.G. Advances in the understanding of the congenital dyserythropoietic anaemias. British Journal of Haematology 131, 431–446 (2005).

4. Swickley, G. et al. Characterization of the interactions between Codanin-1 and C15Orf41, two proteins implicated in congenital dyserythropoietic anemia type I disease. Molecular and Cell Biology 21(2020).

5. Dgany, O. et al. Congenital Dyserythropoietic Anemia Type I Is Caused by Mutations in Codanin-1. The American Journal of Human Genetics 71, 1467–1474 (2002).

6. Renella, R. et al. Codanin-1 mutations in congenital dyserythropoietic anemia type 1 affect HP1α localization in erythroblasts. Blood 117, 6928–6938 (2011).

7. Ask, K. et al. Codanin-1, mutated in the anaemic disease CDAI, regulates Asf1 function in S-phase histone supply. The EMBO Journal 31, 2013–2023 (2012).

8. Babbs, C. et al. Homozygous mutations in a predicted endonuclease are a novel cause of congenital dyserythropoietic anemia type I. Haematologica 98, 1383–1387 (2013).

9. Su, A.I. et al. A gene atlas of the mouse and human protein-encoding transcriptomes. Proceedings of the National Academy of Sciences of the United States of America 101, 6062–6067 (2004).

10. Shroff, M., Knebel, A., Toth, R. & Rouse, J. A complex comprising C15ORF41 and Codanin-1: the products of two genes mutated in congenital dyserythropoietic anaemia type I (CDA-I). Biochemical Journal 477, 1893–1905 (2020).

11. Roy, N.B.A. & Babbs, C. The pathogenesis, diagnosis and management of congenital dyserythropoietic anaemia type I. British Journal of Haematology 185, 436–449 (2019).

12. Schindelin, J., et al. Fiji: an open-source platform for biological-image analysis. Nature Methods 9, 676-682 (2012).

13. Schuck, P. Size-Distribution Analysis of Macromolecules by Sedimentation Velocity Ultracentrifugation and Lamm Equation Modeling. Biophysical Journal 78, 1606–1619 (2000).

14. Manalastas-Cantos, K. et al. ATSAS 3.0: expanded functionality and new tools for small-angle scattering data analysis. Journal of Applied Crystallography 54, 343–355 (2021).

15. Konarev, P.V., Volkov, V.V., Sokolova, A.V., Koch, M.H.J. & Svergun, D.I. PRIMUS: a Windows PC-based system for small-angle scattering data analysis. Journal of Applied Crystallography 36, 1277–1282 (2003).

16. Mirdita, M. et al. ColabFold: making protein folding accessible to all. Nature Methods 19, 679–682 (2022).

17. Svergun, D., Barberato, C. & Koch, M.H.J. CRYSOL - a Program to Evaluate X-ray Solution Scattering of Biological Macromolecules from Atomic Coordinates. Journal of Applied Crystallography 28, 768–773 (1995).

18. Kikhney, A.G., Borges, C.R., Molodenskiy, D.S., Jeffries, C.M. & Svergun, D.I. SASBDB: Towards an automatically curated and validated repository for biological scattering data. Protein Science 29, 66–75 (2020).

19. Kavan, D. & Man, P. MSTools—Web based application for visualization and presentation of HXMS data. International Journal of Mass Spectrometry 302, 53–58 (2011).

20. Schenkmayerova, A. et al. Engineering the protein dynamics of an ancestral luciferase. Nature Communications 12, 3616 (2021).

21. Smit, J.H. et al. Probing Universal Protein Dynamics Using Hydrogen–Deuterium Exchange Mass Spectrometry-Derived Residue-Level Gibbs Free Energy. Analytical Chemistry 93, 12840–12847 (2021).

22. Uhrik, L. et al. Hydrogen deuterium exchange mass spectrometry identifies the dominant paratope in CD20 antigen binding to the NCD1.2 monoclonal antibody. Biochemical Journal 478, 99–120 (2021).

23. Masson, G.R. et al. Recommendations for performing, interpreting and reporting hydrogen deuterium exchange mass spectrometry (HDX-MS) experiments. Nature Methods 16, 595–602 (2019).

24. Perez-Riverol, Y. et al. The PRIDE database and related tools and resources in 2019: improving support for quantification data. Nucleic Acids Research 47, D442–d450 (2019).

25. Micsonai, A. et al. BeStSel: webserver for secondary structure and fold prediction for protein CD spectroscopy. Nucleic Acids Research 50, W90–W98 (2022).

26. Drew, E.D. & Janes, R.W. PDBMD2CD: providing predicted protein circular dichroism spectra from multiple molecular dynamics-generated protein structures. Nucleic Acids Research 48, W17–W24 (2020).

27. Kozeleková, A. et al. Phosphorylated and Phosphomimicking Variants May Differ—A Case Study of 14-3-3 Protein. Frontiers in Chemistry 10(2022).

28. Jumper, J. et al. Highly accurate protein structure prediction with AlphaFold. Nature 596, 583–589 (2021).

29. Ahdash, Z. et al. HDX-MS reveals nucleotide-dependent, anti-correlated opening and closure of SecA and SecY channels of the bacterial translocon. eLife 8, e47402 (2019).

30. Lim, C.J. & Cech, T.R. Shaping human telomeres: from shelterin and CST complexes to telomeric chromatin organization. Nature Reviews Molecular Cell Biology (2021).

31. Veverka, P., Janovic, T. & Hofr, C. Quantitative Biology of Human Shelterin and Telomerase: Searching for the Weakest Point. International Journal of Molecular Sciences 20, 13 (2019).

32. Nandakumar, J. & Cech, T.R. Finding the end: recruitment of telomerase to telomeres. Nat Rev Mol Cell Biol 14, 69–82 (2013).

33. Sedor, S.F. & Shao, S. Mechanism of ASF1 engagement by CDAN1. Nature Communications 16, 2599 (2025).

34. Jeong, T.-K., Frater, R.C.M., Yoon, J., Groth, A. & Song, J.-J. CODANIN-1 sequesters ASF1 by using a histone H3 mimic helix to regulate the histone supply. Nature Communications 16, 2181 (2025).

35. Abramson, J. et al. Accurate structure prediction of biomolecular interactions with AlphaFold 3. Nature 630, 493–500 (2024).

36. Tang, Y. et al. Structure of a human ASF1a–HIRA complex and insights into specificity of histone chaperone complex assembly. Nature Structural & Molecular Biology 13, 921–929 (2006).

37. English, C.M., Adkins, M.W., Carson, J.J., Churchill, M.E.A. & Tyler, J.K. Structural basis for the histone chaperone activity of Asf1. Cell 127, 495–508 (2006).

