## Supplementary_Information for "Inherited CDA-I disease: anemia-associated mutations disrupt CDIN1-Codanin1 complex"

### Supplementary Table

**Supplementary Table 1**: **SEC-SAXS sample, data collection, analysis, and 3D modeling parameters for CDIN1-Codanin1$_{Cterm}$ complex, Codanin1$_{Cterm}$, and CDIN1**[1,2]

| | | | |
|---|---|---|---|
| **(*a*) Sample details** | | | |
| Organism | *Homo sapiens* (Human) | | |
| Source (Catalogue No. or reference) | *E. coli* (BL21) recombinant expression | | |
| | CDIN1-Codanin1$_{Cterm}$ Complex | Codanin1$_{Cterm}$ | CDIN1 |
| *Scattering particle composition* | CDIN1:Codanin1$_{Cterm}$ | Codanin1$_{Cterm}$ | CDIN1 |
| Proteins | CDAN1-interacting nuclease 1 [UniProt Q9Y2V0 (1–281)] with an additional N-terminal SGTLPETGSSEA<br>Codanin-1<br>[UniProt Q8IWY9 (1005–1227)] with an additional N-terminal SG | Codanin-1<br>[UniProt Q8IWY9 (1005–1227)] with an additional N-terminal SG | CDAN1-interacting nuclease 1 [UniProt Q9Y2V0 (1–281)] with an additional N-terminal SGTLPETGSSEA |
| Stoichiometry of components | 1:1 | | |
| *Sample environment/configuration* | | | |
| Solvent composition | 20 mM Tris-HCl, 150 mM NaCl, 5% glycerol, pH 8 | 20 mM Tris-HCl, 150 mM NaCl, 5% glycerol, pH 8 | 20 mM Tris-HCl, 150 mM NaCl, 5% glycerol, pH 8 |
| Sample temperature (°C) | 20°C | 20°C | 20°C |
| In beam sample cell | flow cell | flow cell | flow cell |
| *Size Exclusion Chromatography SEC-SAS* | | | |
| Sample injection concentration (mg ml$^{-1}$) | 10 | 10 | 10 |
| Sample injection volume (ml) | 0.045 | 0.045 | 0.045 |
| SEC column type | Superdex 200 Increase 5/150 GL | Superdex 200 Increase 5/150 GL | Superdex 200 Increase 5/150 GL |
| SEC flowrate, mL/min | 0.3 | 0.3 | 0.3 |
| **(*b*) SAS data collection** | | | |
| Data acquisition/reduction software | | | |
| Source/instrument description or reference | EMBL P12 beamline at the PETRA III storage ring (DESY; Hamburg, Germany) using a Pilatus 6M detector at a sample-detector distance of 3 m | | |
| Measured *q*-range ($q_{min}$ – $q_{max}$, nm$^{-1}$) | 0.0259 – 4.434 | | |
| Method for scaling intensities | Relative scaling | | |
| Exposure time, number of frames. For SEC-SAS, final number of sample frames used for averaging. | 0.495 s/frame (1200 frames)<br>Averaging 9 (712 – 720) | 0.495 s/frame (1200 frames)<br>Averaging 10 (842 – 851) | 0.495 s/frame (1200 frames)<br>Averaging 10 (692 – 701) |
| Additional relevant details | λ = 0.123982 nm | | |

| **(*c*) SAS-derived structural parameters** | | | |
|---|---|---|---|
| Methods/Software | PRIMUS/qt (3.2.1), AUTORG and GNOM | | |
| *Guinier Analysis* | CDIN1-Codanin1$_{Cterm}$ Complex | Codanin1$_{Cterm}$ | CDIN1 |
| $I(0)^{\dagger} \pm \sigma$ (a.u) | 1619.11 ± 3.84 | 1539.76 ± 4.46 | 2018.92 ± 13.22 |
| $R_g^{\dagger\dagger} \pm \sigma$ (nm) | 3.04 ± 0.01 | 2.36 ± 0.01 | 2.81 ± 0.03 |
| *$qR_g$ range (datapoint range)* | 0.57 – 1.26 (57 – 136) | 0.22 – 1.29 (24 – 182) | 0.44 – 1.27 (46 – 148) |
| Linear fit assessment (AUTORG fidelity) | 0 | 0.55 | 0.01 |
| *PDDF/P(r) analysis* | CDIN1-Codanin1$_{Cterm}$ Complex | Codanin1$_{Cterm}$ | CDIN1 |
| $I(0)^{\dagger} \pm \sigma$ (a.u.) | 1612 ± 2.84 | 1533 ± 4.02 | 2022 ± 8.77 |
| $R_g^{\dagger\dagger} \pm \sigma$ (nm) | 3.05 ± 0.01 | 2.36 ± 0.01 | 2.82 ± 0.01 |
| $D_{max}^{\dagger\dagger\dagger}$ (nm) | 10.56 | 7.42 | 8.94 |
| $q$-range (nm$^{-1}$) | 0.1874 – 2.6331 | 0.0663 – 3.1847 | 0.1557 – 2.8429 |

| **(*d*) Scattering particle size** | | | |
|---|---|---|---|
| Methods/Software | $V_p$[1], Volume of correlation $V_c$[3], SAXSMow[4], DatBayes[5] from the ATSAS suite[6] | | |
| | CDIN1-Codanin1$_{Cterm}$ Complex | Codanin1$_{Cterm}$ | CDIN1 |
| *Volume estimates* | | | |
| Porod volume, $V_p$ (Å$^3$) | 101 518 | 46 346 | 104 364 |
| *Molecular weight (M) estimates (Da)* | | | |
| From SAS, concentration independent method | | | |
| SAXS MoW | 55 206 | 26 625 | 62 343 |
| Bayesian inference, range with % confidence | 52 550 – 60 200 (94%) | 26 550 – 29 250 (94%) | 58 800 – 67 900 (92%) |
| $V_p$ | 59 875 | 25 918 | 66 329 |
| $V_c$ | 57 022 | 27 926 | 66 627 |
| Size & Shape | 69 294 | 30 867 | 76 640 |
| From SAS-independent measure (MS, MALS) | n.a., 60 kDa | 24 961 Da, 26 kDa | 33 369 Da, 69 kDa |

| **(*e*) Modelling** | | | |
|---|---|---|---|
| *Atomistic modelling methods* | | | |
| | CDIN1-Codanin1$_{Cterm}$ Complex | Codanin1$_{Cterm}$ | CDIN1 |
| Software | CRYSOL[7] (with default parameters) from PRIMUSqt in ATSAS 3.2.1 | | |
| $q$-range for fit ($q_{min} – q_{max}$; nm$^{-1}$) | 0.075 – 4.434 | 0.072 – 4.434 | 0.012 – 4.434 |
| Symmetry/anisotropy assumptions | no | no | no |
| $\chi^2$, CorMap $P$-values for fit | 6.14, 0.00 | 1.04, 0.04 | 1.06, 0.30 |

| **(*f*) Data and model deposition** | | | |
|---|---|---|---|
| | CDIN1-Codanin1$_{Cterm}$ Complex | Codanin1$_{Cterm}$ | CDIN1 |
| SASBDB IDs | SASDVJ4 | SASDV49 | SASDV39 |

† I(0) – forward scattered intensity; †† $R_g$ – radius of gyration providing an estimated overall protein size; ††† $D_{max}$ – maximum dimension of the particle.

### Supplementary Figures

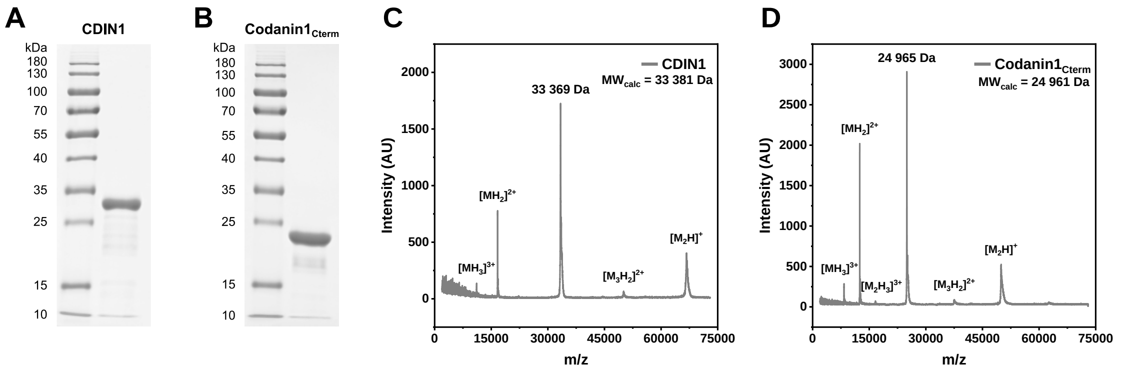


**Supplementary Figure 1.** Proteins' purity and identity analyses. (A, B) Coomassie-stained 12% SDS-PAGE gels show the purity of recombinant proteins CDIN1 and Codanin1$_{Cterm}$ above 95%. CDIN1 (4 µg) and Codanin1$_{Cterm}$ (4 µg) were loaded in corresponding wells after purification. Molecular weight marker PageRuler™ Prestained Protein Ladder (Thermo Fisher Scientific, p# 26616) and band sizes are depicted on the left. (C, D) MALDI-TOF MS analyses of CDIN1 and Codanin1$_{Cterm}$ confirmed the identity of both proteins. The highest observed peaks with experimentally determined molecular weight corresponded to the calculated molecular weights indicated under the proteins' names in plots. The differences between calculated and experimental molecular weights were smaller than the size of a single amino acid, suggesting that the proteins were appropriately prepared, including residues remaining after tag cleavage.

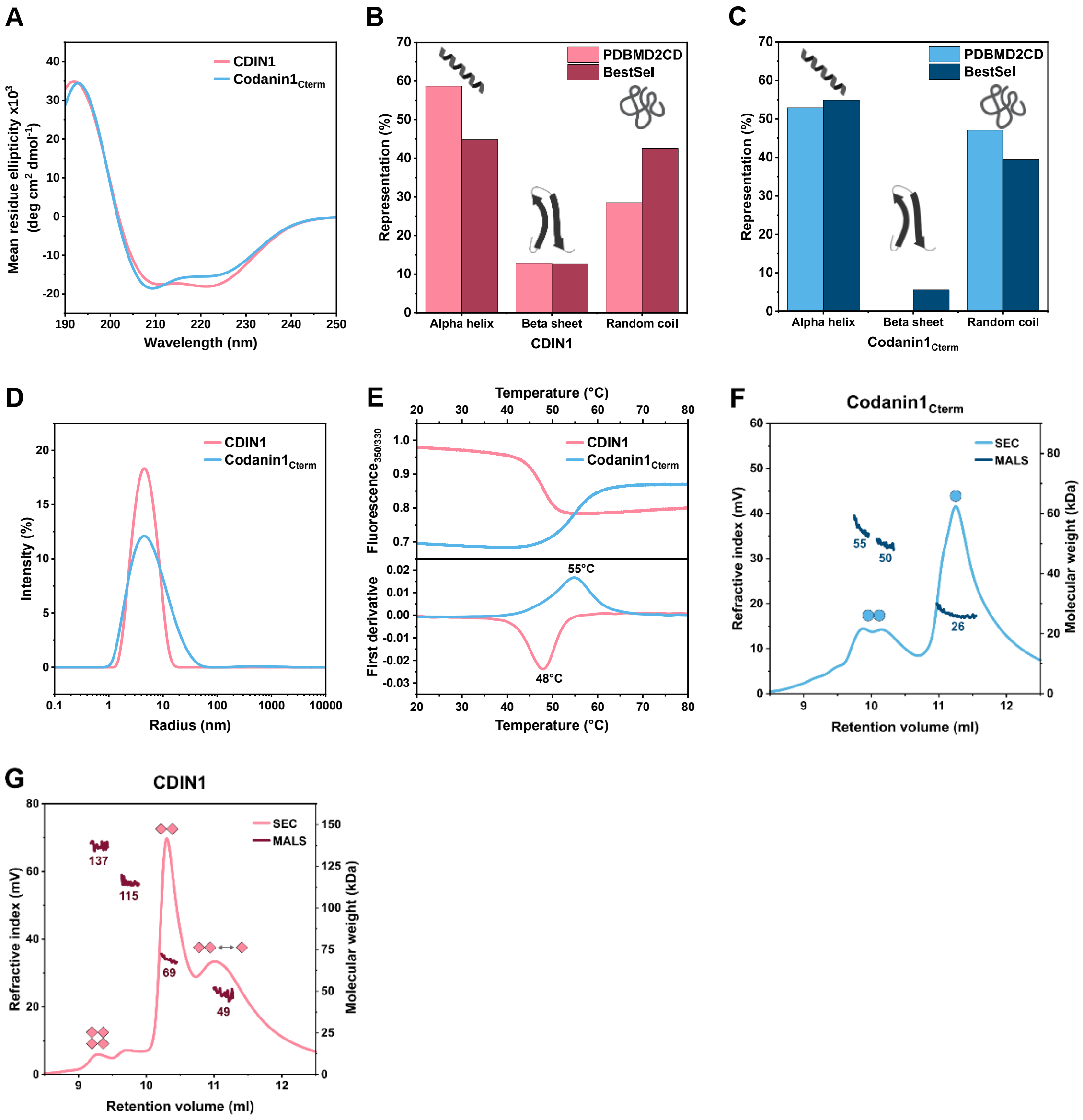


**Supplementary Figure 2.** Secondary structure, homogeneity, and thermal stability of CDIN1 and Codanin1$_{Cterm}$. (A) Far-UV Circular dichroism (CD) spectra of CDIN1 and Codanin1$_{Cterm}$ employed to determine secondary structures' contribution experimentally. (B, C) Bar graphs showing the difference between secondary structure composition calculated from AlphaFold predicted structures using PDBMD2CD software (light color shades) and secondary structure composition determined from experimentally obtained CD spectra analyzed by BeStSel software (dark color shades). (D) The dynamic light scattering (DLS) measurements showing particle radius distribution demonstrate the homogeneity of both proteins, indicating no aggregation. The plot shows the dependence of the percentage of scattering intensity on the particle radius. (E) Differential scanning fluorimetry (nanoDSF) shows thermal stability with inflection points corresponding to 48 °C and 55 °C for CDIN1 and Codanin1$_{Cterm}$, respectively. (F) SEC-MALS showing the dominant monomeric arrangement of Codanin1$_{Cterm}$. (G) SEC-MALS suggests mainly dimeric occurrence of CDIN1. The double arrow represents the dynamic equilibrium between monomeric and dimeric states.

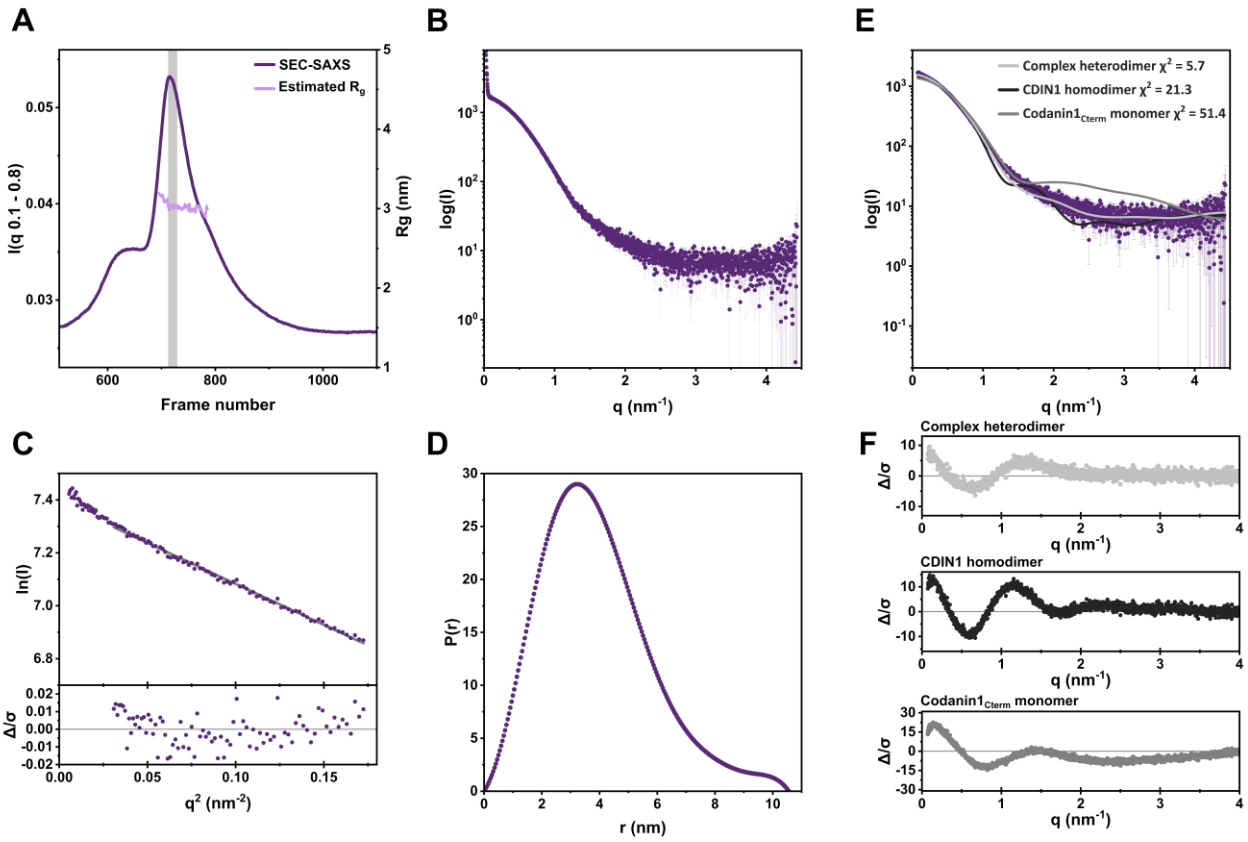


**Supplementary Figure 3.** SEC-SAXS analysis of CDIN1-Codanin1$_{Cterm}$ complex. (A) Estimated $R_g$ (light) and I(q 0.1-0.8) (dark) plots for CDIN1-Codanin1$_{Cterm}$ complex. The gray rectangle highlights frames (712 – 720) used for averaging and further analysis. (B) SAXS intensity profile of CDIN1-Codanin1$_{Cterm}$ complex with subtracted background plotted as log(I) versus q. (C) Guinier plot of the initial part of the scattering curve showing the quality of the measured protein sample. The lower section shows error-weighted residual plot for the linear fit starting at 0.03 $nm^{-2}$ of $q^2$. (D) Pairwise distance distribution function P(r) indicating maximum dimensions $D_{max}$. (E) Fit of the scattering curve calculated from CDIN1-Codanin1$_{Cterm}$ heterodimer AlphaFold model (light gray), CDIN1 homodimer AlphaFold model (black) and Codanin1$_{Cterm}$ monomer AlphaFold model (dark gray) with SAXS data (purple circles) generated by CRYSOL[7]. (F) Error-weighted residual plots for the CRYSOL model fits of CDIN1-Codanin1$_{Cterm}$ heterodimer (light gray), CDIN1 homodimer (black) and Codanin1$_{Cterm}$ monomer (dark gray).

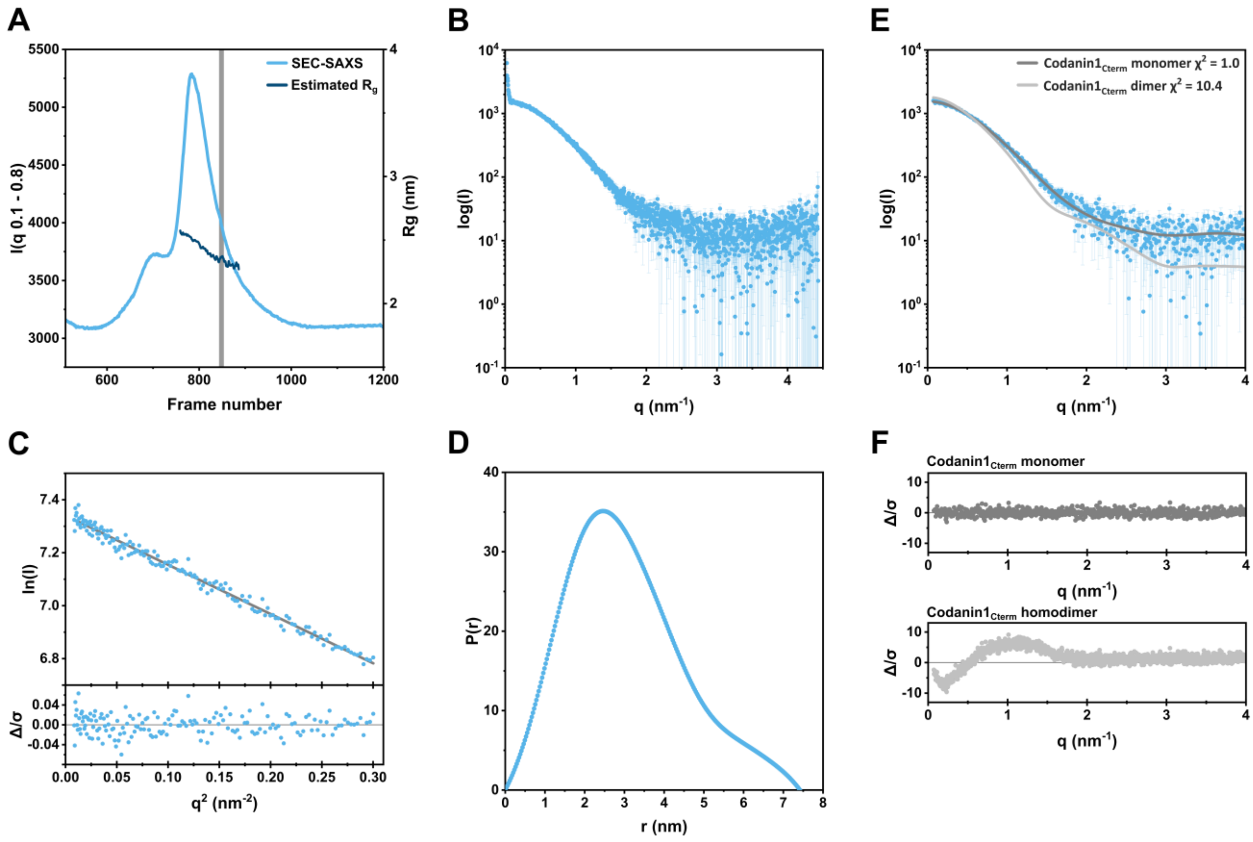


**Supplementary Figure 4.** SEC-SAXS analysis of Codanin1$_{Cterm}$. (A) Estimated $R_g$ (dark) and I(q 0.1-0.8) (light) plots for Codanin1$_{Cterm}$. The gray rectangle highlights frames (842 – 851) used for averaging and further analysis. (B) SAXS intensity profile of Codanin1$_{Cterm}$ with subtracted background plotted as log(I) versus q. (C) Linear Guinier plot of the initial part of the scattering curve showing the quality of the measured protein sample. The lower section shows error-weighted residual plot for the linear fit. (D) Pairwise distance distribution function P(r) indicating maximum dimensions $D_{max}$. (E) Fit of the scattering curve calculated from Codanin1$_{Cterm}$ monomer AlphaFold model (dark gray) and Codanin1$_{Cterm}$ homodimer AlphaFold model (light gray) with SAXS data (blue circles) generated by CRYSOL[7]. (F) Error-weighted residual plots for the CRYSOL model fits of Codanin1$_{Cterm}$ monomer (dark gray) and Codanin1$_{Cterm}$ homodimer (light gray).

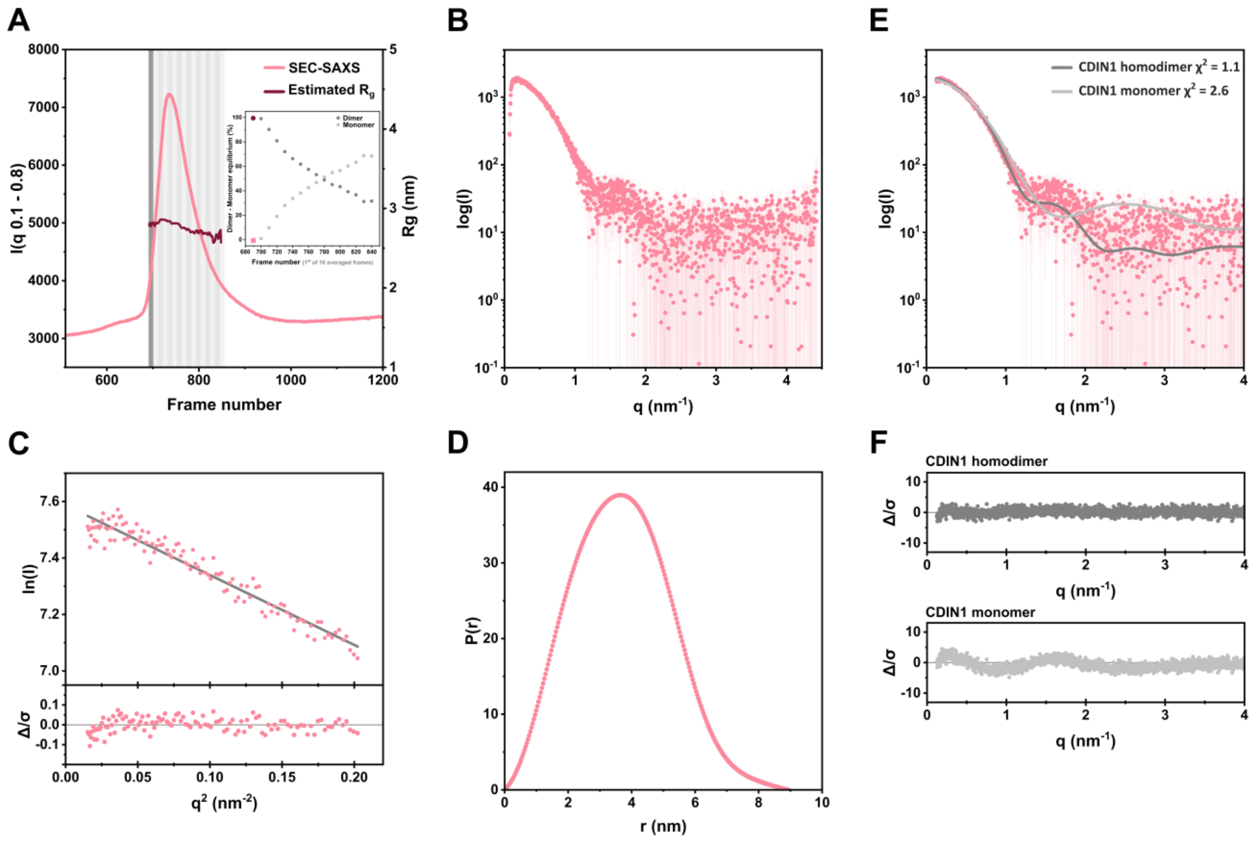


**Supplementary Figure 5.** SEC-SAXS analysis of CDIN1. (A) Estimated $R_g$ (dark red) and I(q 0.1-0.8) (light red) plots for CDIN1. The gray rectangles show frame regions analyzed by OLIGOMER[8] to determine actual contribution of dimer and monomer states. The inlet plot shows the distribution of the dimeric and monomeric state of CDIN1 across frame number regions, divided into intervals of 10 frames. The dark gray rectangle highlights frames (692 – 701) used for averaging and further analysis. (B) SAXS intensity profile of CDIN1 with subtracted background plotted as log(I) versus q. (C) Linear Guinier plot of the initial part of the scattering curve showing the quality of the measured protein sample. The lower section shows error-weighted residual plot for the linear fit. (D) Pairwise distance distribution function P(r) indicating maximum dimensions $D_{max}$. (E) The fit of the scattering curve calculated from CDIN1 dimer AlphaFold model (dark gray) and CDIN1 monomer AlphaFold model (light gray) with SAXS data (pink circles) generated by CRYSOL[7]. (F) Error-weighted residual plots for the CRYSOL model fits of CDIN1 dimer (dark gray) and CDIN1 monomer (light gray).

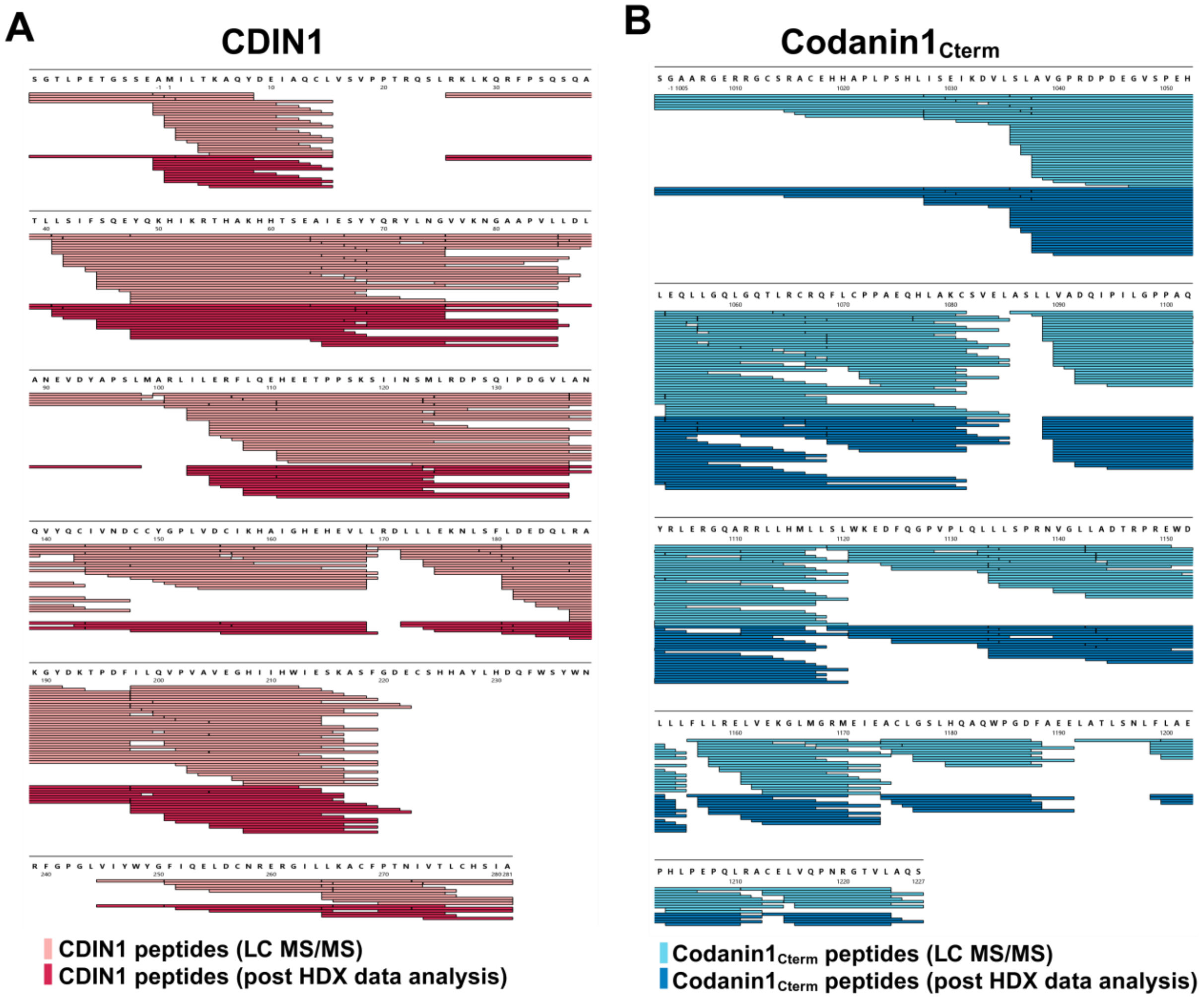


**Supplementary Figure 6.** Proteolytic coverage of CDIN1 and Codanin1$_{Cterm}$ before and after HDX data analysis. Proteolytic peptide mapping of CDIN1 and Codanin1$_{Cterm}$ was carried out using LC-MS/MS under the same conditions as those used for HDX-MS, before and after filtering for high-confidence peptides. Only peptides with a high signal-to-noise ratio and reliable isotopic distribution patterns were retained for HDX analysis. (A) CDIN1 sequence coverage map. A graphical representation of peptide coverage across CDIN1 sequence. LC-MS/MS analysis identified 242 unique peptides (shown as light red bars), covering 89 % of the full CDIN1 sequence. After HDX data evaluation, 108 high-confidence peptides (dark red bars) were retained, covering 87.03 % of the sequence. (B) Codanin1$_{Cterm}$ sequence coverage map. A graphical representation of peptide coverage across the Codanin1$_{Cterm}$ sequence. LC-MS/MS analysis identified 232 peptides (light blue bars), covering 100 % of the full sequence. After HDX data processing, 143 high-confidence peptides (dark blue bars) were retained, covering 95.56 % of the sequence.

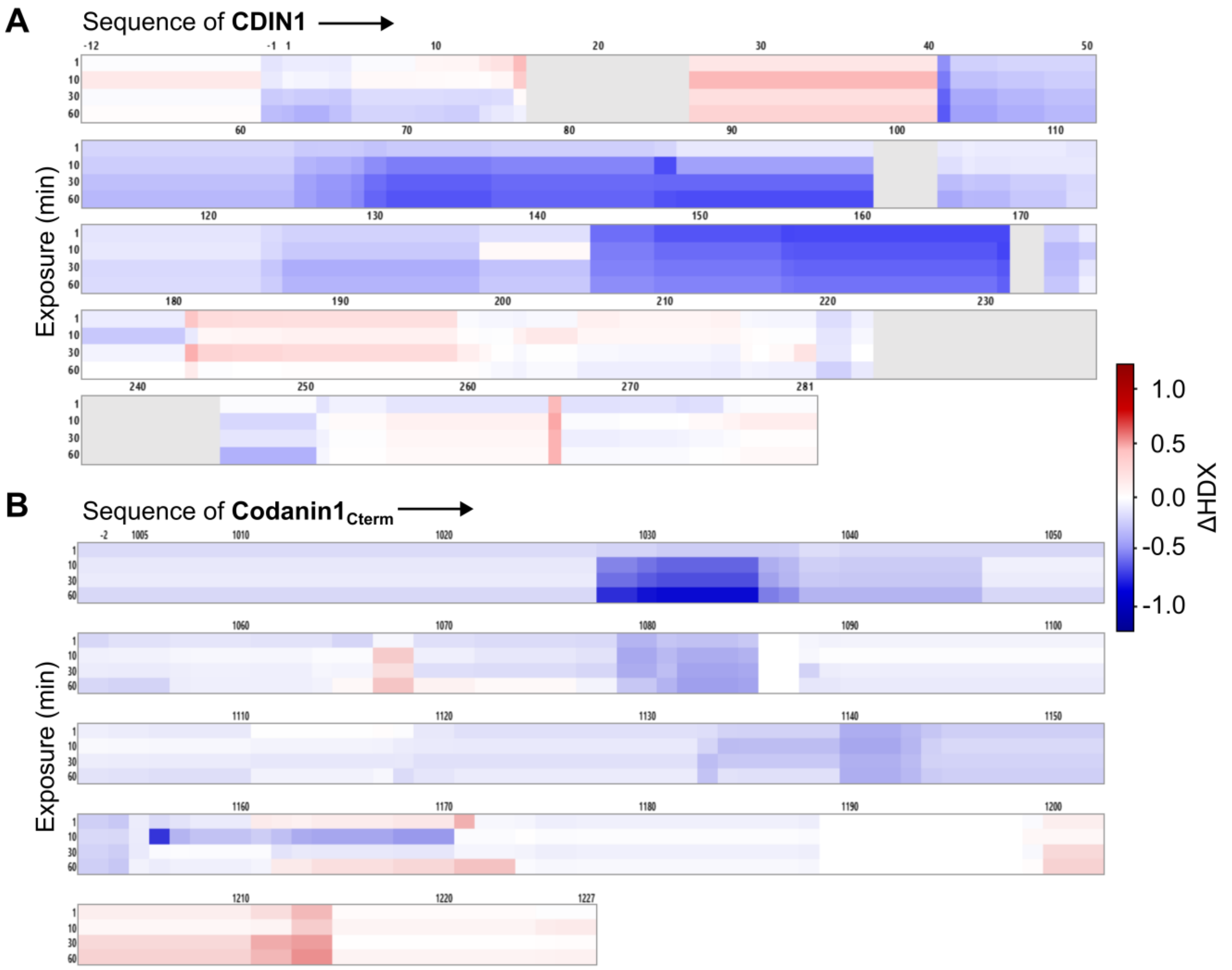


**Supplementary Figure 7.** ΔHDX heatmaps highlighting protein-protein interaction regions identified using HDX-MS. The difference in relative deuterium uptake (ΔHDX) was calculated by subtracting the uptake in the complexed form from that in the free protein (ΔHDX = free protein - complex), thus highlighting changes associated with complex formation. ΔHDX values from all labeling time points (1 min, 10 min, 30 min, and 60 min) were mapped onto the corresponding protein sequences. Negative ΔHDX values (shown in blue) reflect reduced solvent accessibility or increased structural rigidity, suggesting regions involved in binding. Positive ΔHDX values (shown in red) indicate increased solvent accessibility or flexibility in the complex relative to the free form. A calibrated color scale is provided for quantitative interpretation. (A) ΔHDX values for the CDIN1-Codanin1$_{\text{Cterm}}$ complex mapped onto the CDIN1 sequence. Regions with negative ΔHDX values suggest potential interaction sites between CDIN1 and Codanin1$_{\text{Cterm}}$. (B) ΔHDX values for the Codanin1$_{\text{Cterm}}$-CDIN1 complex mapped onto the Codanin1$_{\text{Cterm}}$ sequence. Negative ΔHDX values indicate potential binding sites involved in interaction with CDIN1. Gray regions denote proline residues or sequence segments not covered by HDX-MS analyses.

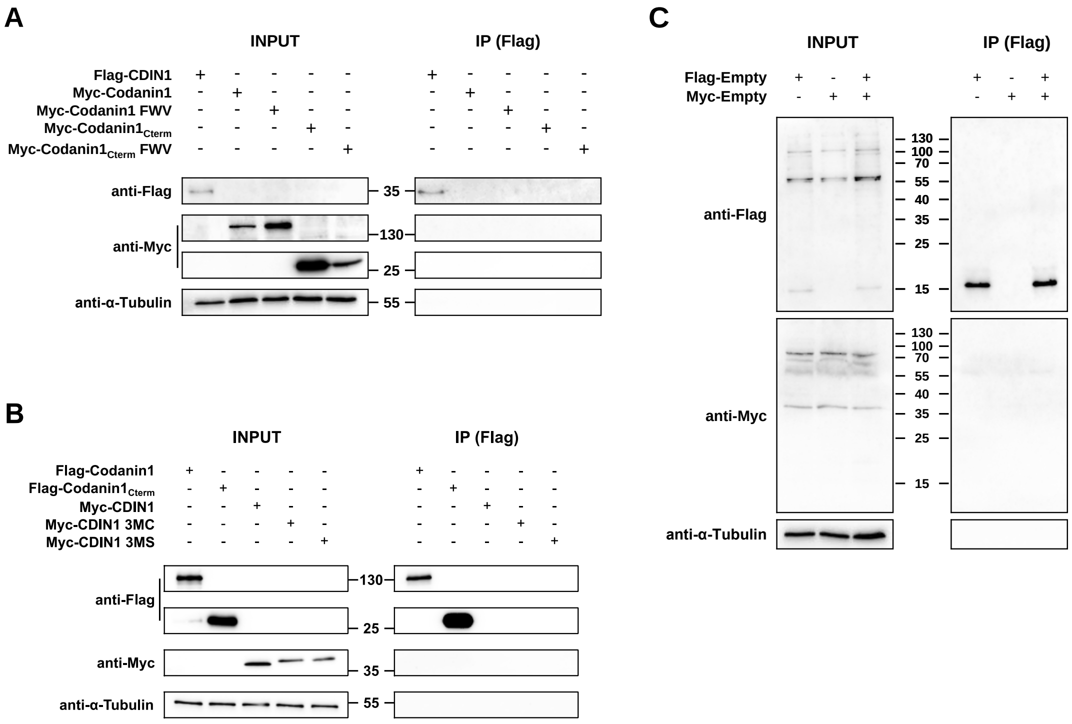


**Supplementary Figure 8.** Co-IP controls confirmed the specificity of antibodies. (A, B) Western blot analyses of Co-IP samples obtained from HEK293T transfected by each of CDIN1 and Codanin1 plasmids separately show the specificity of used antibodies and M2 Flag magnetic beads. (C) Western blot analyses of Co-IP samples obtained from HEK293T transfected by empty vectors detected no non-specific signal at the molecular level of proteins of interest in IP fractions. To ensure the visibility of all recognized fractions, empty vector controls were subjected to an extended exposure time of 120 seconds. In contrast, a short 2-second exposure was sufficient for the detection of Flag- or Myc-tagged proteins, which exhibited a significantly enhanced signal.

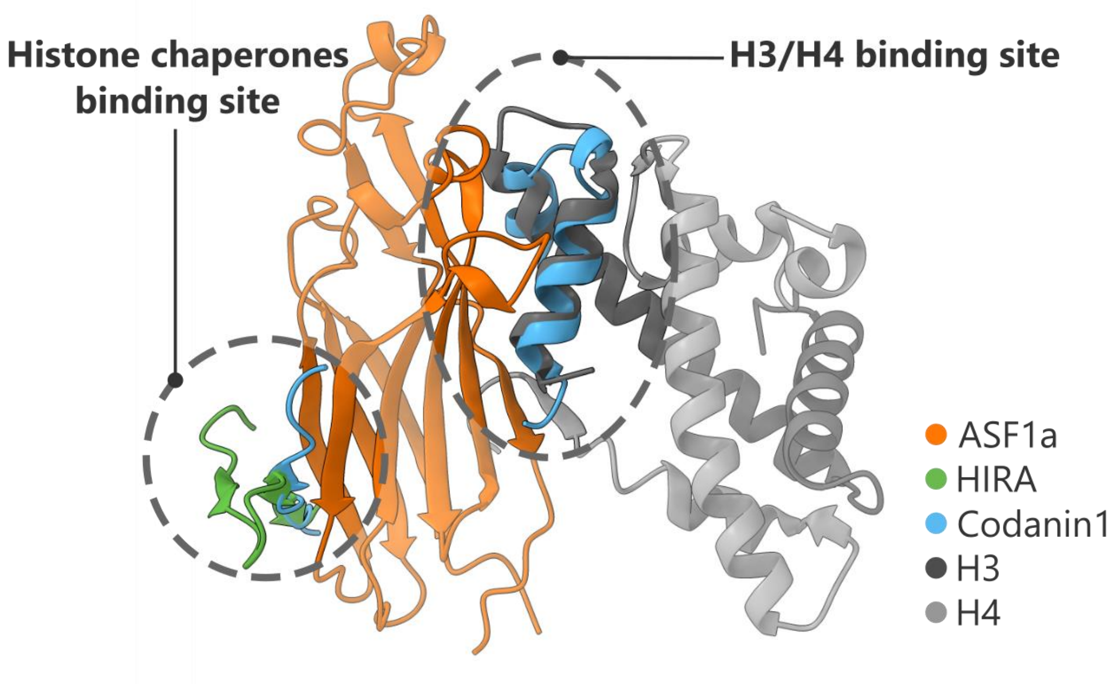


**Supplementary Figure 9**. Overlaid structures demonstrate how human histone chaperone ASF1a interacts with its partners via two binding sites. ASF1a (orange) binds the B-domain of another histone chaperone HIRA (green), within the Histone chaperone binding site. Additionally, ASF1a accommodates H3/H4 dimer (gray) through the H3/H4 binding site. AlphaFold 3[9] predicts that Codanin1 (blue) binds to both ASF1a's interaction sites, as shown by the overlapping Codanin1 structure on ASF1a. Thus, Codanin1 might sterically hinder the interaction with histones and histone chaperone HIRA. Superposition has been produced using published crystallography data ASF1a-HIRA (PDB ID: 2i32)[10], ASF1a-H3/H4 (PDB ID: 2hue)[11], and Codanin1's structure predicted by AlphaFold 3[9] (AlphaFoldDB ID: Q8IWY9).

#### Supplementary References

1. Trewhella, J. et al. 2017 publication guidelines for structural modelling of small-angle scattering data from biomolecules in solution: an update. *Acta Crystallographica Section D* 73, 710-728 (2017).
2. Trewhella, J., Jeffries, C.M. & Whitten, A.E. 2023 update of template tables for reporting biomolecular structural modelling of small-angle scattering data. *Acta Crystallographica Section D* 79, 122-132 (2023).
3. Rambo, R.P. & Tainer, J.A. Accurate assessment of mass, models and resolution by small-angle scattering. *Nature* 496, 477-481 (2013).
4. Piiadov, V., Ares de Araújo, E., Oliveira Neto, M., Craievich, A.F. & Polikarpov, I. SAXSMoW 2.0: Online calculator of the molecular weight of proteins in dilute solution from experimental SAXS data measured on a relative scale. *Protein Science* 28, 454-463 (2019).
5. Hajizadeh, N.R., Franke, D., Jeffries, C.M. & Svergun, D.I. Consensus Bayesian assessment of protein molecular mass from solution X-ray scattering data. *Scientific Reports* 8, 7204 (2018).
6. Manalastas-Cantos, K. et al. ATSAS 3.0: expanded functionality and new tools for small-angle scattering data analysis. *Journal of Applied Crystallography* 54, 343-355 (2021).
7. Svergun, D., Barberato, C. & Koch, M.H.J. CRYSOL - a Program to Evaluate X-ray Solution Scattering of Biological Macromolecules from Atomic Coordinates. *Journal of Applied Crystallography* 28, 768-773 (1995).
8. Konarev, P.V., Volkov, V.V., Sokolova, A.V., Koch, M.H.J. & Svergun, D.I. PRIMUS: a Windows PC-based system for small-angle scattering data analysis. *Journal of Applied Crystallography* 36, 1277-1282 (2003).
9. Abramson, J. et al. Accurate structure prediction of biomolecular interactions with AlphaFold 3. *Nature* 630, 493-500 (2024).
10. Tang, Y. et al. Structure of a human ASF1a–HIRA complex and insights into specificity of histone chaperone complex assembly. *Nature Structural & Molecular Biology* 13, 921-929 (2006).
11. English, C.M., Adkins, M.W., Carson, J.J., Churchill, M.E.A. & Tyler, J.K. Structural basis for the histone chaperone activity of Asf1. *Cell* 127, 495-508 (2006).
